# TRPM8 channels control Golgi morphology through an AR-dependent induction of the Scd1 gene and modulation of unsaturated lipid content

**DOI:** 10.1101/2024.10.14.618203

**Authors:** Mariam Wehbi, Yves Gouriou, Anne-sophie Borowiec, Juliette Geoffray, Sally Badawi, Christophe Chouabe, Christian Slomianny, Dmitri Gordienko, Fabrice Gonnot, Etienne Dewailly, Philippe Delcourt, René Ferrera, Jean-Paul Pais-de-Barros, Mazen Kurdi, Laurent Héliot, Fabien Van Coppenolle, Loic Lemonnier, Natalia Prevarskaya, Gabriel Bidaux

## Abstract

Transient receptor potential melastatin 8 (TRPM8), the cold and menthol receptor is essential to thermosensation, although its roles in organs within the body are still unclear. Besides TRPM8, we previously cloned several isoforms, like 4TM-TRPM8, which can be expressed with or without TRPM8.

In this study, we characterize the human TRPM8(85) in ER membranes in the vicinity of Golgi apparatus (GA) and mitochondria in prostate epithelial cells. Silencing of TRPM8(85) induces lipid droplet accumulation, GA expansion and fragmentation associated with a drop in the vesicular trafficking to plasmalemma. Furthermore, lipidomic analysis reveals a strong shift in unsaturated fatty acids (UFAs), induced by TRPM8(85) silencing and to a lesser extent silencing of TRPM8. UFAs increase is caused by the induction of Δ9 stearoyl desaturase (*Scd1*) gene. Silencing SCD1 or palmitate incubation prevent GA expansion in TRPM8(85)-silenced cells. Finally, we demonstrated that TRPM8 regulates SCD1 via the androgen receptor.

## INTRODUCTION

Originally cloned in prostate cancer (PCa) and later detected in other organs also insulated by the body such as bladder or liver^1,2^ the Transient Receptor Potential Melastatin 8 (TRPM8) has been widely characterized as the cold receptor, whose activation in afferent fibers led to the induction of thermogenic mechanisms and behaviours^3–7^. TRPM8 overexpression has also been associated with melanoma, breast cancer, pancreatic cancer, (for review read^8^). Besides its expression in the neurons of the dorsal root ganglia and cancers, TRPM8 physiological roles have been studied in epidermis^9–11^ and testis^12,13^, immune cells and the digestive system^8^. In internal organs of the body, most studies have focused on the role of TRPM8 in PCa. We and others have characterized 1) TRPM8 as a calcium channel in both plasmalemma and ER membranes of apical epithelial cells in the prostate^14–16^, 2) the regulation of the *trpm8* gene transcription by androgen receptor (AR) and its channel activity by testosterone and AR^17^, and 3) demonstrated that its expression and activity correlated with the PCa grade^16,18–21^. Although the roles of TRPM8 in the cell cycle, cell death, and differentiation of PCa cells have been validated, the molecular mechanisms behind are not fully understood. The mystery surrounding TRPM8 deepens further when considering that the *trpm8* gene does not only gives rise to the canonical cold and menthol receptor but also to a diversity of alternate mRNAs and their related protein isoforms^22^. To facilitate the understanding, we classified the TRPM8 isoforms in 4 groups based on their structure: short non-channel isoforms which are regulatory subunits^23–25^, 4TM-TRPM8 isoforms which are functional channels in ER microdomains^11,26^ and the long isoforms. These latter variants have not been characterized yet. In conclusion, 23 years after the identification of TRPM8, understanding why evolution has selected the cold receptor and its isoforms to be expressed in organs insulated within the body like the prostate, including in homeotherms, remains challenging.

One archetypal cold receptor is the bacterial DesK receptor expressed by Bacillus subtilis. DesK, stands for Desaturase Kinase, is a three-protein system that include a dual phosphatase/kinase cold sensing module in the membrane, a transcription factor and a Δ5 desaturase (for review^27^). Under cold stimulation, the sensing module shifts to kinase activity and activates the Des transcription factor. In turn, the latter is translocated to the nucleus where it trans-activates the Δ5 desaturase gene (Des). Des role is to desaturate the phospholipids in the bacterial membrane what induce a progressive decrease in membrane order - equivalent to decrease in membrane thickness and density^28^. Saturated fatty acids (SFAs) are known to mimic cold by increasing membrane order, whereas mono-unsaturated fatty acids (MUFAs) and PUFAs increase it and thus mimic warmth^27^. In a more disordered membrane, DesK switches back to the phosphatase form and cancels out the activity of the Des transcription factor. It can thus be hypothesized that thermo-sensors, such as DesK, are sensing their lipid environment, and by extension, may function as lipid sensors. Structural and molecular studies revealed that lipids interaction with DesK coiled-coil domain induce its conformational switch^29^ and modulated its thermosensitivity^28^. Noteworthily, the carboxy-terminal loop of TRPM8 ends with a coiled-coil domain downstream at the TRP box, both being involved in temperature sensing^30–32^. Furthermore, TRPM8 has been reported to be directly activated by lipids such as phosphatidylinositol 4,5- bisphosphate^33,34^, lysophospholipids^35^, but inhibited by poly-unsaturated fatty acids, PUFAs^36^. In line with DesK, we wondered whether TRPM8 acts as a lipid sensor that could regulate a feedback loop controlling membrane lipid composition.

In the current study, we used the androgen-resistant prostate metastasis cell line LNCaP C4-2b as a model expressing most TRPM8 isoforms and assessed the shared features in prostate of *wt* and TRPM8 isoforms knock-out (KOM8) mice. We firstly characterized the (9’-26) alternate mRNA and its TRPM8(85) isoforms expressed in both normal and cancer prostate tissue. This isoform is expressed in ER membranes in the vicinity of the Golgi apparatus (GA) and mitochondria but not in the mitochondria-associated membrane like are the 4TM-isoforms. Using differential siRNA-mediated knock-down, we invalidated the different TRPM8 groups and characterized the resulting changes at both the cellular and subcellular levels. We reported that suppressing TRPM8(85) induced lipid droplets accumulation, increased in cell size, a combined expansion and fragmentation of GA and reduced vesicular trafficking between GA and plasmalemma. Lipidomics on TRPM8(85)-KD LNCaP cells and KOM8 prostate revealed a shift in the phosphatidyl Choline (PC) / phosphatidyl Ethanolamine (PE) ratio and the SFAs/MUFAs ratio. This increase in MUFA was driven by an increase in Δ9 desaturase (SCD1) expression following TRPM8(85) KD. Silencing SCD1 or palmitate incubation prevented GA expansion in TRPM8(85)-silenced cells. Finally, we showed that TRPM8 controls SCD1 expression through an AR-dependent mechanism.

## MATERIALS AND METHODS

### Cell lines culture

The LNCaP, DU145 and PC-3 cell lines were purchased from the American Type Culture Collection (ATCC). LNCaP C4-2b cell line was a generous gift from Dr. F. Cabon, Paris. Cells were amplified in RPMI medium 1640 (Gibco®) supplemented with 10% fetal calf serum (FCS) and kanamycin (100 µg/ml). For the experiment, cells were cultured in RPMI medium 1640 (Gibco®) supplemented with 2% FCS, 1mM Sodium pyruvate, 1.5mM CaCl2, and kanamycin (100 µg/ml). LNCaP cells were cultured in RPMI 1640 (1x) + GlutaMAX™ medium supplemented with 10% fetal bovine serum (FBS) and 1% penicillin-streptomycin antibiotic. HEK cell line was grown in Dulbecco’s minimal essential media (DMEM) (Gibco) including 4.5 g/l Glucose and 1.8 mM Ca^2+^ and was supplemented with 10% FCS and Kanamycin (100 µg/ml).

### Viability assay

Cells were transfected with siRNA in 100 mm dishes overnight. The day after, cells were dispatched in 96-well plates at a density of 5,000 cells per well in 100 μl RPMI medium. After 6 hours of incubation, experiments started as day 0, and treatments were applied by adding 100 μl of treatment-containing medium. From day 1, half of the medium was changed daily. CellTiter 96® AQueous Non-Radioactive Cell Proliferation Assay (Promega) was used to determine the number of viable cells each day. **TRPM8^-/-^ Mice**. The establishment of our mouse line has been detailed in a previous study^11^. Briefly, the loxP sites have been inserted between exon 18 and 20 in order to delete the ion pore domain and consequently suppressed the activity of all channel-like TRPM8 isoforms, conversely to other models available worldwide. TRPM8 knock-out (KOM8) mice have backcrossed for several generations in order to obtain a pure C57BL/6J background, so has it been also done for the TRPM8^flox/flox^ (*wt*) littermates. This study was carried out in strict accordance with the law (Articles R214-117 to 121 of the rural code), and the recommendations in the Guide for the Care and Use of Laboratory Animals of the French National Institute for Medical Research (www.recherche-animale.org/). The protocol has been approved by the Committee on the Ethics of Animal Experiments of the University of Lyon 1 (authorization: BH2012-65). All efforts were made to minimize animal suffering.

### Human prostate tissues

Human prostate tissue specimens were obtained from resection surgeries performed on clinical indications in the Urology Department at Hôpital St Philibert (Lille, France), according to the Declaration of Helsinki Principles and the guidelines of the ethical committee of the Medical Center, Hôpital St Philibert prepared in agreement with French law, and informed consent was obtained from all patients. All procedures were approved by the Ethical Committee of Hôpital St Philibert, Université Catholique de Lille (Lille, France). All tissue specimens came from patients who had not received any anti-androgen therapy. In addition, all resection specimens have been diagnosed by an anatomopathological examination. After patient surgery, the conjunctive tissues were eliminated and the epithelial nodules were cut into small fragments and frozen in liquid nitrogen for storage.

### Transfection

Plasmid transfection for the TRPM8 isoform was performed using X-tremeGENE HP DNA Transfection Reagent (Roche) at a dose of 4 µg per million cells. DharmaFECT (cat#T-2010-03) was utilized for siRNA transfection, including 52.6 nM siRNA for immunoblotting, 62.5 nM siRNA for Golgi expansion assessment and for transfecting 2 µg of plasmid to overexpress SCD1 in LNCaP cells or to express the androgen receptor 640x (1µg). The siRNA sense sequences were as follows: 5’- CUGCAGAAUGGAGGAGAUA(dTdT)-3’ targeting SCD1, 5’-UCUCUGAGCGCACUAUUCA(dTdT)-3’ targeting exon 7 of TRPM8 (siM8-7), 5’-GGGAUGAAAUUGUGAGCAA(dTdT)-3’ targeting exon 10 of TRPM8 (siM8-10), 5’-GGGAUGAAA*<u>G</u>*UGUGA*<u>A</u>*CAA(dTdT)-3’ targeting exon 10 of TRPM8 but with 2 mismatches (siM8-10^mut^), 5’-GGAAACUGGUUGCGAACUU(dTdT)-3’ targeting exon 12 of TRPM8 (siM8-12) and 5’-UAUUCCGUUCGGUCAUCUA dTdT-3’ targeting exon 20 of TRPM8 (siM8-20). For siCTL, negative control siRNA are comprised of 19 complementary RNA bases having no significant homology to any known mouse, rat or human sequences.

### RT-PCR

In brief, total RNA was isolated from different cell lines with TRI Reagent® (SIGMA). Human samples were resuspended in TRI Reagent® (SIGMA) and subjected to two cycles of 30 s at 6,500 × g in a high-throughput PRECELLYS24 tissue homogenizer cooled down with CRYOLIS at 4 °C. After the TRI Reagent extraction has been completed, samples were treated with DNase I (Life Technologies) and reverse transcribed with MuLV reverse transcriptase (Perkin Elmer). PCR was performed using 1 U AmpliTaq Gold (Perkin Elmer). The whole procedure has been described elsewhere^22^. PCR primers are listed in **supplemental table 1**.

### Quantitative real-time PCR analysis

After total mRNA extraction and purification with TRI REAGENT^®^ (Sigma-Aldrich), mRNA samples were incubated with 0.25 µl DNAse (Ambion) per µg of RNA for 25 min at 25°C. Afterwards, mRNA were mixed in a 25:24:1 (V/V) phenol/chloroform/AIA solution (Fluka) with 5% sodium acetate 3 M, incubated for 20 min on ice and finally centrifuged at 15,000 g for 10 min. The upper phase was transferred into a clean 1.5 ml tube supplemented with 10% (volume) of sodium acetate 3M and 25% (volume) of 100% ethanol. After vortexing, tubes were incubated at -20°C overnight in order to precipitate mRNA. Following a brief wash in 70% ethanol, the obtained pellets were left to dry and then re- suspended in 30 µl water. After the quality of mRNA samples was verified on 1% agarose gel, 2 µg of mRNA samples were reverse transcribed. Real-time quantitative PCR has been performed on the Cfx C1000 system (Biorad). For each reaction, the 15 μl final mixture contained 12.5 ng of cDNA, 7.5 µl of 2x SsoFast™ EvaGreen^®^ Supermix (Biorad) and 200 nM primer pairs (see **supplemental table 1**). The housekeeping gene glyceraldehyde-3-phosphate dehydrogenase (GAPDH) was used as an endogenous control to normalize gene expression with the comparative ΔΔCt method. The PCR protocol consisted of initial 30 sec denaturation step at 95° C followed by 40 cycles each comprising 4 sec at 95° C, 30 sec at 60° C and monitoring of final dissociation curve to control the specificity of the amplification.

### Cloning of human SCD1 mRNA

Human prostate mRNA (3µg) have been reverse transcribed with PrimeScript Reverse Transcriptase (Takara). Specific primers (see **supplemental table 1**) targeting the new 5’-extremties of TRPM8 mRNAs have been used to amplify Human prostate cDNA with Phusion Hot Start II High-Fidelity DNA Polymerase (Finnzymes; Thermo Fisher Scientific, Waltham, MA). After a 0.8% (w/vol) agarose-gel extraction of specific DNA bands (Wizard SV gel and PCR Clean-Up System; Promega, Madison, WI), PCR products and recipient vector pVLL-42-SYFP2^37^ were digested with High Fidelity *Sac*I and *Not*I (New England Biolabs, Ipswich, MA) at 37°C overnight, cleaned-up on column prior to be ligated with T4 ligase (New England Biolabs) at 16 °C overnight. Products were transformed in JM109 chemo-competent bacteria (New England Biolabs). Plasmid clones were extracted and sequenced before experiments.

**Cloning of human TRPM8 mRNAs and HA-tagged fusion proteins** has been described elsewhere^22^. **Subcellular fractionation** has been described elsewhere^26^. Briefly, 10 75cm^2^-flasks of LNCaP C4-2b cells have been mechanically disrupted with a glass potter homogenizer in 5 ml of solution B, and submitted to several centrifugation to separate whole cells, the post-nuclear fraction and microsomes. The resulting pellet has been suspended in 1 ml of solution B prior to be loaded on 10%:30% Iodixanol gradient prepared with OptiPrep^™^ Density Gradient Medium (SIGMA). Tubes were centrifuged on a SV40 rotor at 100,000 ***g***, 4°C for 2h. Four visible rings of biological matter were divided in four fractions (from 1 to 2 ml), completed with 6 ml of HEPES buffer and subjected to a second ultracentrifugation on a 50Ti rotor at 100,000 ***g***, 4°C for 1h. All pellets isolated from the whole procedure have been either solubilized in an ice-cold buffer (as described in immunoblotting section, below) and sonicated prior to perform immunoblotting.

### Immunoblotting

An ice-cold buffer (pH 7.2) containing 10 mM PO4Na2/K buffer, 150 mM NaCl, 1 g/100 ml sodium deoxycholate, 1% Triton X-100, 1% NP40, a mixture of protease inhibitors (Sigma-Aldrich), and a phosphatase inhibitor (sodium orthovanadate; Sigma-Aldrich) was applied to previously PBS-washed cells in dishes. After 30 min incubation on ice, the protein extract was transferred into 1.5 ml tubes and sonicated. After 10 minutes of centrifugation at 14,000 g, the pellet was transferred into a clean tube prior to a determination of the protein concentration using a BCA Protein Assay (Pierce). Proteins were either denaturated at 60°C for 30min (for TRPM8 detection) or at 95°C for 5 minutes. An SDS-page was performed using 25-50 µg of total protein loaded into a 7.5% (for the detection of TRPM8, IP3R1 and Giantin) and 10% (for other proteins) polyacrylamide gel. After electrophoresis, proteins were transferred to a PVDF membranes (for TRPM8 detection) or a nitrocellulose membrane using a semi-dry electroblotter (Bio-Rad). The membrane was blocked in a TNT +5% (W/V) milk (15 mM Tris buffer, pH 8, 140 mM NaCl, 0.05% Tween 20, and 5% non-fat dried milk) for 30 min at room temperature, then soaked in the primary antibody diluted in TNT +1% milk for either 2 h at room temperature or overnight at +4°C. After three washes in TNT, the membrane was soaked in the secondary antibody diluted in TNT+1% milk for 1h at room temperature. The membrane was processed for chemiluminescence detection using Luminata Forte Western HRP Substrate (Millipore) according to the manufacturer’s instructions. After a 10 min bath in Re- blot PLus Mild solution (Millipore), membrane was blotted again. The primary antibodies were: rabbit anti- TRPM8 (Ab109308, Abcam), rabbit anti-HA tag (Sc-805, Santa Cruz), goat anti-GAPDH (Sc-20357, Santa Cruz), mouse anti-SCD1 (ab19862; 1/1000) and mouse anti-GAPDH (Sc- 47724; 1/1000).

### ImmunocytoFluorescence

Experiments were performed on LNCaP C4-2b cells plated on 35 mm glass bottom dishes (MatTek Inc). LNCaP C-42b cells were fixed with 4% formalin in PBS for 10 min on ice prior to 3 PBS washes. Cells/tumors slices were subjected to blocking and permeabilization with PBS + 1.2% gelatine + 0.2% Tween + 0.2M glycine for 30 min at 37°C. The slides/dishes were then incubated with primary antibodies 2 h at 37°C. After thorough rinsing in PBS/gelatine, the slides/dishes were treated with the corresponding secondary antibody: either Dye light 488-labeled anti-rabbit IgG (Jackson ImmunoResearch; dilution, 1/2000) or Alexa fluor 546-labeled anti-mouse IgG (Molecular Probes; dilution, 1/4000) diluted in PBS/gelatine for 1h at ambient temperature. After rinsing twice in PBS/gelatine and once in PBS with 1/200 Dapi for 10 min at ambient temperature, the slides were mounted with Mowiol® and examined under a confocal microscope. The primary antibodies used were: rabbit anti-TRPM8 (Ab109308, Abcam), mouse anti-p21waf1 (Clone SX118, Dako), Anti-Giantin antibody [9B6] - cis/medial Golgi Marker (ab37266, Abcam), Anti-GM130 antibody [EP892Y] - cis-Golgi Marker (ab52649, Abcam), Anti-Golgin-97 antibody- trans-Golgi Marker (ab84340, Abcam), Anti-LMAN1 antibody [EPR6979] - ERGIC marker (ab125006, Abcam), Anti-HA tag antibody - ChIP Grade (ab9110, Abcam), Anti-SCD1 antibody [CD.E10] (ab19862, Abcam). For assessment of GA expansion, LNCaP cells were plated on 8-well glass slide (ref PEZGS0816). After 48 or 96 hours of siRNA transfection, cells were washed once with PBS 1X then fixed using 4% paraformaldehyde for 10 minutes. After fixation, cells were washed 3 times with PBS 1X then permeabilized for 5 minutes using 0.1% Triton/PBS. Blocking was done using 1% BSA for 45 minutes. Then primary antibody (anti mouse-Golgin-97: Abcam, ab169287) was incubated overnight at 4°C at dilution 1/1000. After 3 washes with PBS 1X, secondary antibodies were incubated in dark at room temperature: Alexa Fluor 647-labelled anti-mouse IgG (Invitrogen A21235) or Alexa Fluor 488-lablled anti- mouse IgG (Invitrogen A11001); dilution, 1/1000. After 30 minutes, cells were washed 3 times with PBS 1X then the slides were mounted using Fluoromount^TM^ aqueous mounting medium and topped with cover glass before examination under the confocal microscope.

### Flow cytometry

Cells were harvested, split into 1 million cell samples in 15 ml tubes before fixation with 1 ml of 70% Ethanol at −20°C overnight. Cells were washed twice with PBS/4% BSA 0,1% Triton ×100 and finally incubated at RT for 30 min. Cell were then pelleted by centrifugation at 250 × g for 8 min at 20°C. For analysis of immunolabeled cell population, mouse anti- p21 (Clone SX118, Dako) primary antibody was diluted in 100μl PBS-BT at 1/200 and incubated with cells at ambient temperature for 1 h. After a first quick wash in PBS, a second wash was done at RT for 30 min. Anti-mouse IgG coupled to Alexa Fluor 647 (Jackson ImmunoResearch; dilution 1/4000) was then incubated at RT for 30 min. After two PBS washouts for a total incubation time of 30 min at ambient temperature, cells were suspended in 500μl of PBS and then analyzed. Flow cytometry was performed with a CyAnTM ADP Analyser. Data were analysed with FlowJo software (version 8.7).

### Viability Assay

Cells were transfected with siRNA in 100 mm dishes overnight. The day after, cells were dispatched in 96-well plates at a density of 5,000 cells per well in 100 µl RPMI medium. After 6 hours of incubation, experiments started as day 0, and treatments were applied by adding 100 µl of treatment-containing medium. From day 1, half of the medium was changed daily. CellTiter 96^®^ AQueous Non-Radioactive Cell Proliferation Assay (Promega) was used to determine the number of viable cells each day.

### Confocal imaging

Immunofluorescence imaging, VSVG-GFP assay, Laurdan experiments were done on a Nikon Eclipse Ti, A1R confocal microscope with a 40x and 60x oil objective and equipped with the following lasers: diode 405 nm (blue), argon ion 488-514 nm (blue-green), diode 561 nm (orange-red) and diode 642 nm (deep red). Multiple filter settings were used according to the needs. Static images were acquired by averaging 4-scanning lines for noise reduction and better resolution. Pixel size was set around 70-100nm per pixel. Spectral detection for Laurdan experiments was done using a 32 channels GASP on a bandpass 400 to 600nm.

For live cell imaging experiments on the Nikon confocal microscopy experiments the cells were bathed in physiological salt solution (PSS) containing (in mM): NaCl 140, KCl 5, MgCl2 1, glucose 10, HEPES 10; pH adjusted to 7.4 with NaOH. PSS was supplemented with 3 mM EGTA or 2 mM CaCl2, depending on experimental protocol, as described in the text.

### Image analysis

For colocalization analysis, confocal 8-bit images were filtered to remove signal below the mean fluorescence intensity of the image prior to be analysed by the module Coloc 2 of Image J2 (v2.14.0). Mander’s correlation was chosen to obtain the colocalization value of each channel. Original image had a pixel size between 70-100nm per pixel in order to respect Shannon criterion and get the maximal resolution. For Golgi expansion assessment, Golgi surface area of each cell was measured using the plugin 3D objects counter of ImageJ (Version 1.54k 15 September 2024). The macro selected the regions of interest corresponding to the GA structure for identified objects larger than 20 pixels and the surface area was measured by conversion of pixel dimension.

### Membrane order measurement

Membrane order was measured in LNCaP C4-2b cells after 48 hours of siRNA transfection, SCD1 overexpression or 3 days after palmitate and oleate (50µM each) treatment. For this, 10µM of laurdan probe (6-Dodecanoyl-2-Dimethylaminonaphthalene) (Thermofisher, Cat no. D250) was added to the cells in RPMI 1640 (1x) + GlutaMAX™ lacking FBS for 45 minutes in incubator set at 37°C, 5%CO2 then washed with PBS 1X. Before acquisition, cells were fixed using 4% paraformaldehyde (PFA) for 10 minutes in the incubator set at 37°C, 5% CO2. After fixation, cells were washed with PBS 1X 3 times. Image acquisition was achieved on a confocal microscopy with a spectral APD and a 60x objective. Generalized Polarization (GP) of laurdan was calculated as: GP=(I440-I490)/(I440+I490)^38^ with a home-made macro (available on demand) running under Image J and the distribution of image pixels by their GP value was calculated for each image. An average distribution was then calculated from a minimum of 5 images for each of the three days of experiments.

### VSVG-GFP assay

This protocol includes double transfection and it was done in 35 mm imaging dishes with glass bottom (cat. No: 81168). LNCaP C4-2b cells were seeded and at the end of the day, transfection with different siRNAs was done. After 24 hours of siRNA transfection, transfection was done with 0.5µg DNA encoding the mutated vesicular stomatitis virus G protein fused in frame with the green fluorescent protein (VSVG-GFP) using DharmaFECT (cat # T-2010-03) solution. After 24 hours of VSVG transfection, cells were incubated at the restrictive temperature of 40°C with a humidified atmosphere of 5% CO2 at 37°C for 12 hours. Cells were then transferred to permissive temperature at 32°C on the microscope to initiate the trafficking of VSVG and record image time-series at a rate of 1 image per minute. Images were analysed on the setup with NIS-Element. Transport time were determined as following:

- ER to GA: was determined when fluorescence out of the GA reached a minimum.
- GA to Vesicle budding: was determined by the time interval between the time at which fluorescence out of the GA reached a minimum and the time at which vesicle budding started to become obvious.
- Vesicle budding to plasmalemma was determine by the time interval between obvious figures of vesicle budding and first stretch of fluorescence on the cell border.

Only cells with a bright intensity were included in the analysis. Values were calculated for each cell. At least 10 images were analysed for each of the three days of experiments.

### Cytosolic Ca^2+^ imaging

Living cells were imaged on an inverted epifluorescence microscope Leica DMi6000B using a 40x oil-immersion objective, with Lambda DG4 wavelength-switch xenon light source (Sutter Instruments), equipped with an ORCA-Flash4.0 digital CMOS camera C11440 (Hamamatsu). Cytosolic Ca^2+^ level was measured using ratiometric dye fura-2 (2 µM) and quantified according to the Grynkiewicz equation^39^. The bath solution (HBSS (Hank’s Balanced Salt Solution) contained 142 mM NaCl, 5.6 mM KCl, 1 mM MgCl2, 2 mM CaCl2, 0.34 mM Na2HPO4, 0.44 mM KH2PO4, 10 mM HEPES and 5.6 mM glucose. The osmolarity and pH of external solutions were adjusted to 310 mOsm.l^−1^ and 7.4, respectively. The cells were continuously perfused with the HBSS solution and chemicals were added *via* a perfusion system.

### Mitochondrial Ca^2+^ imaging

Living cells were imaged on an Axiovert 200M inverted microscope attached to an LSM 510 META laser-scanning unit (Zeiss) and running LSM 510 software (Zeiss). Mitochondrial Ca^2+^ level was estimated by imaging the fluorescent Ca^2+^ indicator rhod-2. 50 μg of rhod-2-AM were dissolved in 10 μL of DMSO [containing 0.025% (wt/vol) pluronic F-127], which was then mixed with 4 mL of PSS and superfused to the experimental chamber for 20 min. The incubation of the cells with the dyes was followed by a 1-h wash in PSS containing 1.7 mM CaCl2 to allow time for de-esterification. The dye loading was performed at RT. The cells were then kept for 30 min at 37 °C. Before imaging was commenced the cells were superfused with PSS containing 70 μM CaCl2 and supplemented with 10 μM LaCl3 to eliminate capacitative Ca^2+^ entry. The cells were stimulated with 200 μM menthol. Rhod-2 fluorescence intensity (F) was normalized to the average fluorescence intensity in the images acquired before agonist application (F0). The temporal profiles of the agonist-induced [Ca2+]m transients are illustrated by the plots showing the time course of the normalized rhod-2 fluorescence intensity (F/F0) averaged within a confocal optical slice of the cell.

### Time domain-fluorescence lifetime imaging microscopy for FRET measurements (FRET-FLIM)

For live-cell imaging, cells were placed on 35mm glass bottom dishes (MatTek Corporation, USA), filled with L-15 medium without phenol red (Life technologies), and kept at 37 °C using a stage incubator (Life Imaging Services, Switzerland). FLIM was performed with a Leica TCS SP5 X confocal head (Leica Microsystems, Germany) with the SMD upgrade, mounted on an inverted microscope (DMI6000, Leica Microsystems, Germany). A pulsed diode laser, PDL 800-B (PicoQuant GMBH, Germany), delivered 40 MHz repetitive rate pulses at 405 nm. The confocal pinhole was set to 1 Airy, for a 0.921 μm optical slice. Single photon events originated from the illuminated voxel were collected through a 63×/1.2 NA water-immersion objective and recorded by a TCSPC detector (HydraHarp 400; PicoQuant GMBH, Germany). Fluorescence was detected through a 483/Δ32 single-bandpass filter (Semrock, USA) on Single Photon Avalanche Photodiodes, SPAD (MPD, Italy), set up at 256 × 256 pixels. Single photon arrival times were measured with SymPhoTime software (PicoQuant GMBH, Germany) while image were computed with LAS AF software (Leica Microsystems, Germany. In order to obtain the best resolution of organelles, a 5-fold zoom factor was applied, giving a pixel size of 0.193 μm and an image size of 49.21 × 49.21 μm. Since the statistical analysis of the distribution of single photon arrival times requires at least 100 photons per pixel, 120 frames were acquired at 200 Hz and integrated in the final image. Fluorescence lifetime of the fluorescence donor was determined by the Phasor plot method using a custom-developed software [19]. Since fluorescence lifetime is independent of the fluorescence emitter concentration, possible cell-to-cell variations of the fluorophore concentration can be neglected in the FRET-FLIM measurements. FRET efficiency value, namely E(FRET), was calculated according to the equation: E(FRET)= 1-(**τ**_DA_/**τ**_D_). Note that E(FRET) is directly proportional to the fraction of Ca^2+^- bound Cameleon, and, hence, to steady-state [Ca^2+^].

### High-pressure Freezing, Freeze Substitution (HPF/FS) and Immuno-Electron Microscopy

Cells were scratched and collected by centrifugation at 200g for 5 min. The cell pellet was gently suspended in a low volume of 20% BSA as a cryoprotective agent. The mixture was transferred into the 3 mm specimens carriers (Leica Microsystems Inc, Vienna, Austria), and then was high-pressure frozen with a Leica EM HPM100 High Pressure Freezer (Leica Microsystems Inc, Vienna, Austria) at -197°C, 210 KPa and maintained under liquid nitrogen.

Samples were then cryo-substituted in a Leica AFS2 system (Leica Microsystems, Vienna, Austria) prior to be transferred under liquid nitrogen to freeze substitution medium consisting of methanol and PAF 0.5%. Samples were incubated at -85°C for 1 hour 30 minutes, and then were warmed to -45°C over a period of 14 hours. Prior to embedding in K4M resin the samples were washed three times with 100% methanol. Polymerization was performed under UV light for 36 hr. Ultrathin sections (90nm) were obtained as described above.

For immuno-electron microscopy, sections were incubated in blocking medium (0.2 M glycin, 5% donkey serum, 0.2% BSA-c^TM^, 0.1% fish gelatin in PBS buffer) for 30 minutes and then were exposed to primary antibodies: 1/50 anti-calnexin (MAB3126, Millipore) and 1/30 anti-HA (715500, Invitrogen) overnight at room temperature. After washing, the sections were incubated at room temperature for 2 hours in the corresponding secondary gold-conjugated antibodies (Jackson ImmunoResearch) diluted in PBS + 1.2% gelatin. Following thorough wash in PBS and then in water, the sections were stained with 0.5% uranyl acetate for 12 min. Imaging was performed on Hitachi H600 transmission electron microscope at 75 kV accelerating voltage.

### Morphology analysis by Transmission Electron Microscopy

For morphology analysis, cells were fixed in 2.5% glutaraldehyde dissolved in 0.1 M cacodylate buffer and were post-fixed in 1% osmium tetroxide in the same buffer. After acetonitril dehydration, the pellets were embedded in Epon. Serial thin sections (90 nm) were cut using a Leica UC7 ultramicrotome and collected on 150 mesh hexagonal barred copper grids. After staining with 2% uranyl acetate prepared in 50% ethanol and incubation with a lead citrate solution (Reynolds), sections were observed on a Hitachi H-600 transmission electron microscope at 75kV.

### Electrophysiological recordings of TRPM8 channel activity

For GUVs Patch-clamp experiments, we used Axopatch 200B amplifier and pClamp 10.0 software (Molecular Devices, Union City, CA) for data acquisition and analysis. Patch pipettes were fabricated from borosilicate glass capillaries (World Precision Instr., Inc., Sarasota, FL) on horizontal puller (Sutter Instruments Co., Novato, CA) and had a resistance in the range of 7-10 MΩ. Prepared vesicles were immersed in a bath solution containing, in mM, 150 NaCl, 5 glucose, 10 Hepes, pH 7.2. In order to prevent activity of IP3R and RyR channels, commonly present in ER, this solution was supplemented by 1 mM MgCl2, 1 mM EGTA and 1 μM dantrolene (Nelson et al., 1996). Patch pipettes were filled with the same solution, supplemented by 2 μM PiP2 and 1 mM MgATP in order to enhance TRPM8 activity (Rohacs et al., 2005).

### Cell capacitance measurements

Using the whole-cell configuration of the patch-clamp technique, membrane capacitance recordings were made in voltage clamp at room temperature (22-25°C) in LNCaP C4-2b cells 72-96 hours after being transfected with siCTL, siM8-7, siM8-10 or siM8-20. Patch pipettes had resistances of 2-3 MΩ when filled with the internal solution containing (in mM): KCl 140, MgCl2 1, CaCl2 2.5, EGTA 4 and HEPES 10 (pH adjusted to 7.2 with KOH). The external solution contained (in mM): NaCl 140, KCl 5, CaCl2 2, MgCl2 1, Na2HPO4 0.3, KH2PO4 0.4, NaHCO3 4, glucose 5 and HEPES 10 (pH adjusted to 7.4 with NaOH). Membrane capacitance was measured and calculated by analysing the capacitive surge produced by a small voltage step as described previously^40^.

### Lipidomics analysis on LNCaP cells

Chemicals of the highest grade available were from Sigma Aldrich (Saint-Quentin Fallavier, France). LCMSMS quality grade solvents were purchased from Fischer Scientific (Illkirch, France).

Cells (2 million in 200 µl of saline) were spiked with an internal standard mix containing 4 µg of C21:0, 10 µg of epicoprostanol, 0.1 µg of d18:1/17:0 Ceramide, 0.25 µg of d18:1/17:0 sphingomyelin, 1 µg of (14:0)2- PC, 0.1 µg 19:0 LysoPC, 0.2 µg of (14:0)2-PE , 0.1 µg 14:0 Lyso PE and 0.4 µg (14:0)2-PS in a final volume of 90 µl of chloroform/methanol 2/1 (Sigma Aldrich, France). Lipids were extracted according to the method of Folch with 4410 µl of chloroform/methanol 2/1 for 1 h followed by 1250 µl of saline for 1 h at room temperature. Phospholipids and ceramides were analysed by LCMS² as previously described^41,42^.

Total cholesterol and total fatty acids were analysed by GCMS according to Blondelle et al.^43^ Fatty acid quantification was carried out by calculating the response of each fatty acid relative to the 21:0 used as an internal standard.

### Lipidomic analysis on mouse prostates

#### Samples crushing

Samples of tissue were mixed with saline to 0,2 and 0,1 mg tissue/µl. They were further crushed in an Omni Bead Ruptor 24 apparatus (Omni International, Kennesaw, USA) with circa ten 1,4 mm OD zirconium oxide beads (S=6.95 m/s, T=30s, C = 3; D = 10s). Former homogenates (50 µl) were tenfold diluted with saline and crushed again for the extraction of phospholipids and total fatty acids analysis.

*Targeted LCMS² quantification of sphingosines.* Tissue homogenates (5 mg of tissue) were spiked with d17:1 S1P (15 ng) used as internal standard and processed as previously described^44^ with modifications: samples were extracted overnight at 48°C with 1ml of methanol (M) and 0.5 ml chloroform (C). After centrifugation pellets were re-extracted with 1 ml of M/C 2/1 and. Organic phases were combined and evaporated to dryness under vacuum. Dried extracts were finally solubilized with methanol 40 µl. The UHPLC-MS/MS analysis was performed using a Vanquish binary LC system coupled with an Altis plus triple quadrupole mass spectrometer (ThermoElectron).

Separation was achieved at 50 °C on an Eclipse C8 2.1x100 mm, 1.8 µm column (Agilent) with a mobile phase consisting of Solvent A (water containing 0.2% formic acid and 1mM ammonium formate) and Solvent B (methanol containing 0.2% formic acid and 1mM ammonium formate). The gradient elution started at 25% Solvent A and 75% Solvent B for 1 min, transitioning linearly to 100% Solvent B over 5 minutes, held for 5 minutes, and then re-equilibrated back to the starting conditions. The flow rate was set at 0.300 mL/min and an injection volume of 2 μL.

The mass spectrometer was equipped with a H-ESI source, using settings of Ion Voltage at 3500 V, Sheath Gas at 50, Aux Gas at 10, Ion Transfer Tube temperature and Vaporizer temperature at 150°C and 250 °C respectively. Acquisitions were realized using the positive SRM mode: d17:1 P-choline (451.3 → 184, CE 18V), d17:1 Sphingosine-1P (366.3 → 250.2, CE 16V), d18:0 Sphinganine (302.2 → 266.2, CE 16V), d18:0 Sphinganine-1P (382.2 → 284.2, CE 12V), d18:1 P-choline (465.3 → 184, CE 18V), d18:1 Sphingosine (300.2 → 264.2, CE 16V), d18:1 Sphingosine-1P (380.3 → 264.2, CE 16V), d18:2 Sphingosine-1P (378.3 → 262.2, CE 16V). Relative quantitation of sphingosines was performed by calculating the response ratio of the considered molecule to d17:1S1P used as internal standard.

#### Targeted LCMS² quantification of phospholipids and ceramides

An internal standard mix containing the following standards (ng/µl) was prepared : (21:0)2 PC (20, used for PI quantification), d18:1/12:0 SM (25), (19:0)2 PC (50), (14:0)2 PE (25), (17:0)2 PS (20), d18:1/12:0 Cer (2.57), d18:0/12:0 DHCer (0.5), d18:1/12:0 HexCer (2.57), d18:1/12:0 LacCer (0.57) (Sigma Aldrich, France).

Tissue homogenates (equivalent to 2 mg of tissue) were completed to 200 µl of saline and were further spiked with 10 µl and 20 µl of SI-mix respectively.

Lipids were extracted according to the acidified Bligh and Dyer extraction method^45^. Phospholipids and ceramides were analysed by LCMS² as previously described^41,42^. <u>Targeted GCMS² quantification of total fatty acids:</u> Tissue homogenates (equivalent to 0,25 mg tissue) were processed as previously described^43^.

### Data analysis

Data processing and statistical analyses were conducted with Prism 9.0 (GraphPad) software. Before proceeding to any analysis, the normality of the samples was evaluated (Kolmogorov- Smirnov test). Unpaired *t*-test (for normal distribution) or Mann–Whitney test (for non-normal distribution) was used for pairwise comparison unless stated otherwise in the figure legends. One-way ANOVA, Kruskal Wallis test or Two-way ANOVA were used for multiple comparison. For lipidomics analysis, pairewise t-test are given for indication since the statistical power is too weak for most of the lipid species measured. Principal component analysis has been used to reduce the dimension ad sort out the main features responsabile for the variance in lipid contents. Data show mean with standard deviations (SD) or median and 95% confidence intervals (95IC) calculated from at least three independent experiments. For single-cell imaging analysis, statistics were performed on *n* = number of cells to assess single-cell effect as well as heterogeneity between them. Both number of cell and of days of experiments are indicated in the figure legends. A *p* value < 0.05 was considered significant.

## RESULTS

### TRPM8(85) isoform a primate specific isoform at the ER-GA interface in epithelial prostate cells

We previously cloned 33 alternate TRPM8 mRNA in human tissues^22^. TRPM8(9’-26) mRNA was predicted to code for a 85kDa TRPM8 channel form lacking the first 380 amino acids (**Fig. 1A**). TRPM8(9’- 26) mRNA was expressed at a higher level than canonical TRPM8 mRNA and 4TM-TRPM8 mRNAs in PCa cell lines and total human prostate (**Fig. 1B**). In addition, the expression of TRPM8(9’-26) mRNA was independent of the pathological status of the human prostate (**Fig. 1C**). A recombinant HA-tagged TRPM8(85) plasmid expressed in HEK cells confirmed the identification of 85kDa protein (**Fig. 1D**). Immunoblotting of endogenous TRPM8 in LNCaP C4-2b cells detected several proteins at about 120, 95, 85, 75, 55, 45 kDa which were consistent with the detection of the canonical TRPM8, TRPM8(85) and 4TM- TRPM8^10,11,22,26^ (**Fig. 1E**). Interestingly, although TRPM8(9’-26) mRNA had the highest expression level among the TRPM8 mRNAs tested, its related protein showed a lower level than the other isoforms. As demonstrated previously, human alternate TRPM8 exon 9’ is not conserved in rodents, which explains why no 85kDa protein could be detected in the mouse prostate. Surprisingly, the canonical TRPM8 protein was not detected, however higher molecular weight proteins (above 140 kDa) and a doublet around 45kDa corresponding to 4TM-TRPM8s were observed (**Fig. 1F**). All these bands were absent in the prostate of KOM8 mice. These results confirmed that TRPM8(85) did not exist in mouse prostate conversely to 4TM- TRPM8.

**FIGURE 1.**
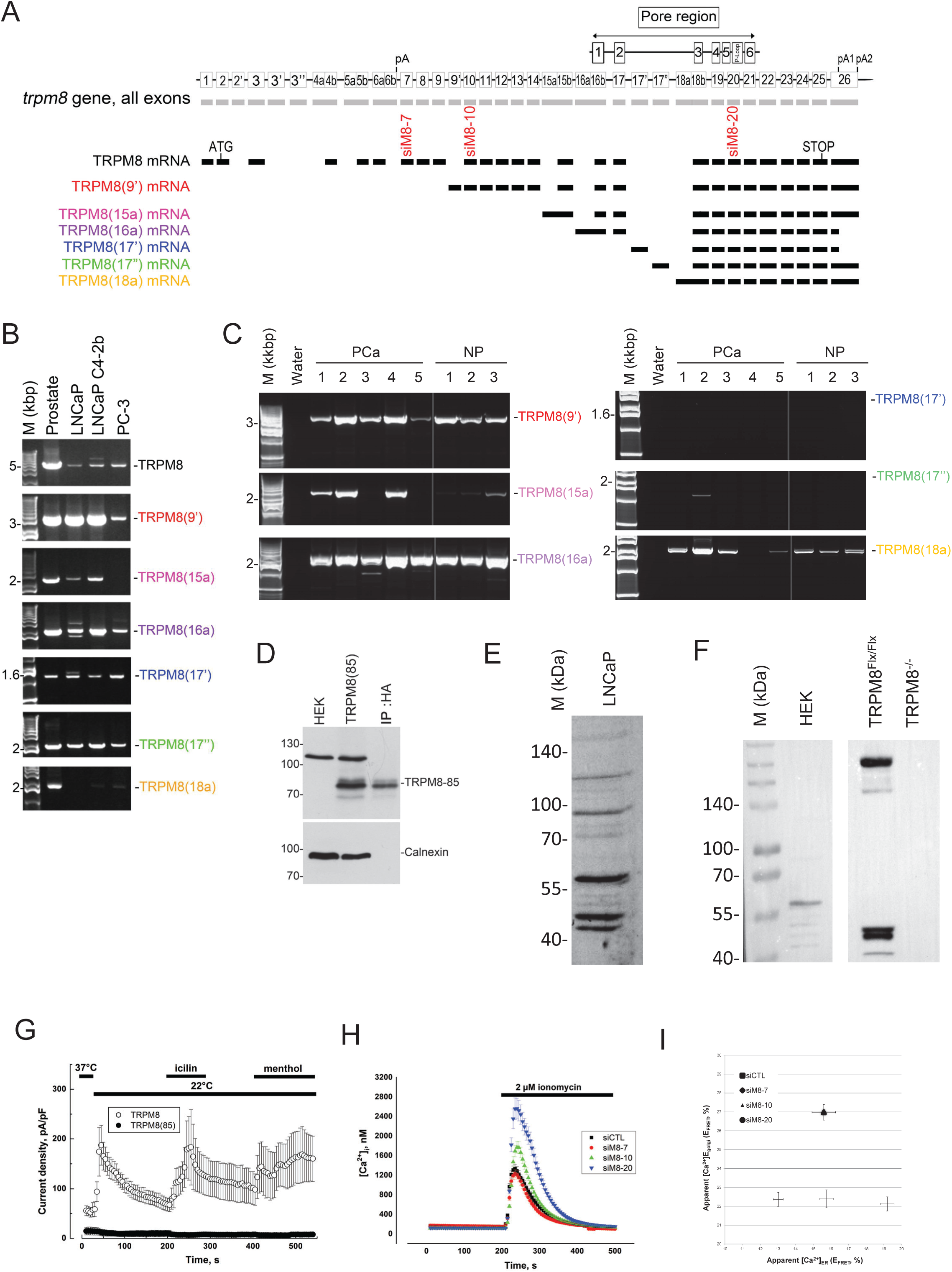
A 85 kDa TRPM8 isoform participates to the Ca^2+^ homeostasis in the Golgi apparatus (GA) of epithelial cells of human prostate. **A,** Schematic representation of the trpm8 gene with all exons. Canonical TRPM8 mRNA (black) displayed with the core exons of trpm8 gene. Below are represented the exon structure of alternate TRPM8 mRNAs: TRPM8(15a), TRPM8(16a), TRPM8(17’), TRPM8(17’’) and TRPM8(18a). Transmembrane (TM) domains and p-loop of the channel pore (Pore region) are aligned with their encoding exons. Position of the different siRNA targeting TRPM8 mRNAs and used in this study is depicted by label "siM8-exon number" below their respective exons. Scheme has been adapted from^11^. **B,** Full length amplification of TRPM8 cDNAs by PCR in normal human prostate tissue (Prostate), androgen- dependent prostate cancer cell line (LNCaP), androgen-refractory prostate cancer cell line (LNCaP C4-2b), androgen-independent prostate cancer cell line (PC-3). **C,** Full length amplification of TRPM8 cDNAs by PCR in different human samples of prostate cancer (PCa) and normal prostate (NP). **D,** Immunoblotting shows the detection of a 85 kDa band in protein samples from HEK cells expressing HA-tagged TRPM8(85) but not in non-transfected cells. Right lane: immunodetection of the HA-tagged TRPM8(85) protein with an anti-TRPM8 antibody after its immunoprecipitation with an anti-HA antibody. Calnexin shows equal protein loading. **E,** Western blot with an anti-TRPM8 antibody on a total protein extract from LNCaP cells showing the different TRPM8 protein sizes presenting the isoforms (repeated three time independently). **F,** Western blots on HEK cells (negative control) and on prostate tissues taken from *wt* (TRPM8^Flox/Flox^) and TRPM8 KO (TRPM8^-/-^) mice (independent tissues analysed: 3 for each group). **G,** Whole-cell patch clamp recordings were conducted in full length TRPM8- or TRPM8(85)-transfected HEK cells. The cell membrane potential was repetitively altered by voltage ramps from−100 mV to +100 mV (applied at 0.2 Hz). The averaged changes in the mean current density at +100 mV, elicited by the exposure to cold (22 °C), 10 μM icilin, 500 μM menthol are plotted on the graph. n=20. **H,** Time course of Ca^2+^ imaging experiments realized on LNCaP C4-2b cells 3 days after transfection with 50 nM of siRNA. Silencing the targets of interest was achieved with the transfection of control siRNA (siCTL; *n* = 387), siRNA targeting full length TRPM8 (siM8-7; *n* = 348), co-targeting TRPM8 and TRPM8(85) (siM8-10; *n* = 345) or co-targeting TRPM8, TRPM8(85) and 4TM-TRPM8 (siM8-20; *n* = 345). Experiments were carried out in the absence of extracellular calcium. **I,** The steady-state Ca^2+^ concentrations in the ER ([Ca^2+^]_ER_) and Golgi apparatus ([Ca^2+^]_Golgi_) were assessed by measuring FRET efficacy through fluorescence lifetime quantification (FRET-FLIM) of fluorescent biosensors in LNCaP C4-2b cells. Cells were co-transfected with plasmids encoding Cameleon biosensors targeted either to ER (D1ER) or to the Golgi apparatus (GoD1CPV) for 24h and a siRNA for 3 days (like in H panel). Time-correlated photon counting (TCSPC) technic was chosen to limit the artefacts caused by variation in biosensor concentration in the organelles. Plots compare the mean lifetime of Cameleon fluorescence in GA (top) and ER (bottom). Values depicted are mean±SD of 30 cumulated cells in 3 independent experiments. After checking the normality of τ_mean_ distribution, an unpaired t-test was performed to compare siM8 with siCTL distributions.

Next, we assessed whether TRPM8(85) could function as an ion channel. No current was detected at the plasmalemma by patch-clamp, suggesting a lack of translocation from the ER to the plasma membrane, which was expected since the first 200 amino-terminal amino acids were lacking^46^. Calcium imaging in HEK cells overexpressing TRPM8(85)-HA showed a very transient calcium increase in the cytosol in very few cells, making it impossible to conclude a stable channel activity (**Fig. S1A**). This could be explained by small concentration of active channels (likely regarding the low level of protein expression) and/or a restricted expression to small Ca^2+^ stores or microdomains. No significant Ca^2+^ increase could be detected in the mitochondria of HEK cells overexpressing TRPM8(85)-HA, conversely to 4TM-TRPM8 overexpression (**Fig. S1B**). We thus looked for an indirect proof of TRPM8(85) activity through the modification of Ca^2+^ stores. We thus quantified the total free Ca^2+^ content in LNCaP cells subjected to pooled silencing of TRPM8 isoforms as presented in previous studies^13,24,26^. As reported in the **figure 1H**, total free Ca^2+^ content was unchanged when only the canonical TRPM8 was silenced (siM8-7) but it increased significantly when TRPM8 and TRPM8(85) were silenced concomitantly by siM8-10. Grouped suppression of TRPM8, TRPM8(85) and 4TM-TRPM8 drastically increased the total free Ca^2+^ content in LNCaP C4-2b cells (siM8- 20). We thus inferred that the endogenous activity of TRPM8(85) had a significant but smaller effect on Ca^2+^ homeostasis than the one of 4TM-TRPM8. Measurements of steady-state Ca^2+^ concentration by FRET-FLIM in the ER and the trans-Golgi network (TGN) by means of D1ER and Go-D1cpv^47^ sensors showed that group suppression of TRPM8, TRPM8(85) and 4TM-TRPM8 increased ER Ca^2+^ content (**Fig. 1I and Fig. S1C**). Surprisingly, co-silencing of TRPM8 and TRPM8(85) was associated with an increase of Ca^2+^ concentration in the TGN. Finally, the suppression of the sole TRPM8 form had no effect on the GA but slightly increase the Ca^2+^ concentration in the ER similar to the one observed under the co-silencing of TRPM8, TRPM8(85). According to previous studies, these results suggested that the canonical TRPM8 form exerted a small activity in the ER membrane^14,15^ while 4TM-TRPM8 has a stronger endogenous activity in ER membranes^26^. Remarkably, we found a hint that TRPM8(85) expression could regulate the Ca^2+^ homeostasis in the TGN but only in the presence of 4TM-TRPM8. Interestingly, modifications in the Ca^2+^ concentration in the TGN has already been reported to cause changes in Golgi apparatus (GA) morphology^47^.

We next assessed the location of TRPM8(85) by immunofluorescence in LNCaP C4-2b cells. We found that both overexpressed HA-tagged TRPM8(85) and endogenous TRPM8s colocalized strongly with an ER marker (**Fig. 2A** and **2B**, respectively). HA-tagged TRPM8(85) partially colocalized with the ERGIC- 53 protein (**Fig. 2C**), a specific marker of the ER–Golgi intermediate compartment (ERGIC) and with Golgin-97 (**Fig. S2A**), a marker of TGN, and with mitochondria (**Fig. 2D**). However, no colocalization was observed with endosomes (**Fig. S2B**). We interpreted these partial colocalizations between ER-seated TRPM8(85) and either ERGIC-53, Golgin-97 or the mitochondrial marker as a close proximity between the ERGIC, TGN and the mitochondrial compartments. Subcellular fractionation and western-blots confirmed that endogenous TRPM8(85) was detected in the post-nuclear fraction (which includes ER membranes), microsomes and migrated with intermediate fractions between the heaviest fractions containing mitochondria and ERGIC and the lightest fractions containing GA in LNCaP C4-2b cells (**Fig. 2E**). Noteworthily, TRPM8(85) segregated in the same fraction as IP3R1 channels while 4TM-TRPM8 segregated with the ERGIC-53 and VDAC fractions. Note that siM8-10 transfection silenced efficiently TRPM8(85) expression in the different fractions.

**FIGURE 2.**
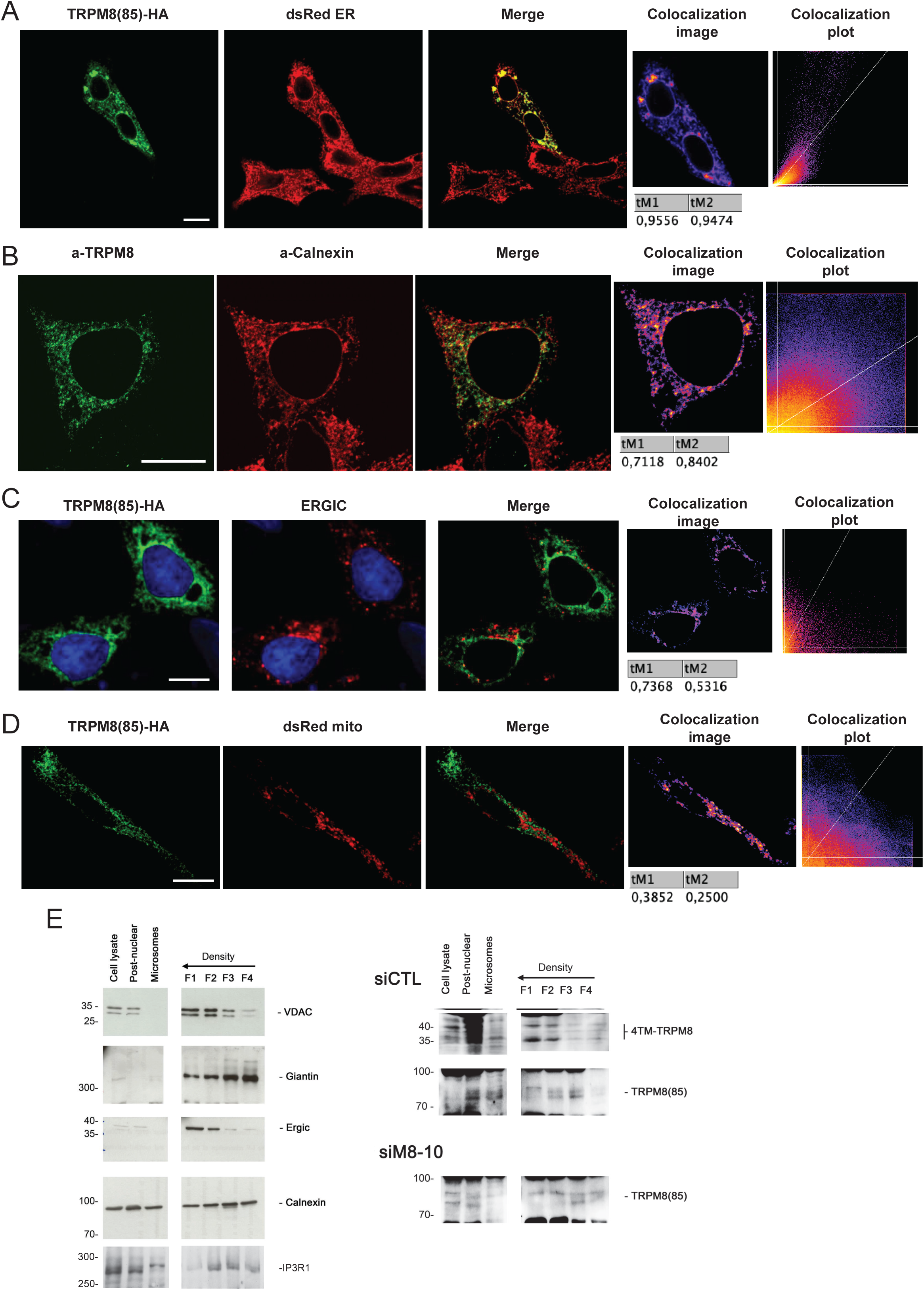
TRPM8(85) is expressed in ER membranes, partly in the vicinity of ERGIC, GA and mitochondrial compartments. Immunofluorescences (**A, B, C and D**) show, in green (1^st^ column), the detection of recombinant HA-tagged TRPM8(85) protein or native TRPM8 channels in LNCaP C4-2b cells, in red (2^nd^ column) ER/ERGIC or mitochondrial markers and overlay of the two fluorescent signals is shown in the 3^rd^ column (Merge). On the last column, colocalization image, colocalization plot and the results of the colocalization pixel analysis with Manders’ coefficient analysed with ImageJ are presented. Scale bar: 10µm. Experiments were reproduced three times independently. **A,** Recombinant HA-tagged TRPM8(85) protein in green with recombinant dsRed targeted to ER in red. **B,** native TRPM8 channels in green with native ER calnexin in red. **C,** Recombinant HA-tagged TRPM8(85) protein in green with ERGIC53 labelling the ERGIC compartment in red. **D,** Recombinant HA-tagged TRPM8(85) protein in green with recombinant dsRed targeted to mitochondria in red. **E,** Cell fractionation and iodixanol density gradient separation (Fraction F1-4) separate organelles of siCTL transfected LNCaP C4-2b cells (siCTL) and demonstrated a partial overlap in TRPM8(85) and 4TM-TRPM8 localization. Calnexin (ER marker), Giantin (cis-GA marker) voltage-dependent anion channel, VDAC (mitochondrial marker) and ERGIC53 (ERGIC marker) The heaviest fractions (F1 and F2) is primarily formed by mitochondrial, associated ER membranes and ERGIC, while the lightest fraction (F4) is mainly composed of Golgi membranes. Fraction F3 is mostly deprived in mitochondria, MAMs, ERGIC but show both strong cis-GA and ER markers. The protein load was 80 µg per fraction. Specificity of TRPM8(85) detection is assessed by its knock-down in siM8-10 transfected cells (siM8-10). Proteins above 80 kDa were separated with 8% gels while proteins lower mass have been separated with 14% gels. Images of immunoblots were sliced to highlight the lanes of interest and the results from several gels were combined in the final figure in order to simplify the reading.

Altogether these results showed that TRPM8(85) isoform is expressed in ER membranes of the human prostate, but not in the mouse, and that it contributes to Ca^2+^ homeostasis in the TGN likely by possibly regulating other TRPM8 rather than through direct ion channel activity.

### TRPM8 and TRPM8(85) co-silencing induces GA expansion, lipid droplets accumulation and slows down vesicles export to plasmalemma

By means of our grouped suppression strategy targeting TRPM8 isoforms, we analysed phenotypic changes in LNCaP C4-2 cells in order to identify possible mechanisms involving TRPM8(85) isoform. Gene expression of ER stress markers (**Fig. S3A**), mitochondrial fusion/fission and stress (**Fig. S3B**) and cell cycle (**Fig. S3C**) revealed that co-silencing of TRPM8 and TRPM8(85) decreased *PCNA* gene expression while it increased *Cdkn1a* gene expression (P21Cip1/Waf1 protein). Additional suppression of the 4TM-TRPM8 (**Fig. S3C**) reverted the regulation of *PCNA*, did not change *Cdkn1a* expression but increased *Cdkn1b* gene expression (P27Kip1 protein). The increase in P21 protein expression was confirmed by flow cytometry in both siM8-10 and siM8-20 groups (**Fig. S3D**). Cell growth of LNCaP C4-2 cells was measured over 4 days after siRNA transfection and we found that only the co-suppression of TRPM8 and TRPM8(85) reduced by about 10% the doubling time (**Fig. S3E**). Compared with the silencing of short TRPM8 isoforms, which exerted a very strong cytostatic effect^24^, we hypothesized that the small anti-proliferative effect of the TRPM8 and TRPM8(85) co-silencing was a side-effect of another mechanism.

We next investigated the subcellular morphological changes by TEM, especially in the GA. Conversely to siCTL group, siM8-7 and siM8-20, siM8-10 induced two major modifications: an expansion of the GA with numerous vesicles and accumulation of lipid droplets (**Fig. 3A and S4**). Since the accumulation of lipid droplets or vesicles could be expected to increase cell size, we evaluated the latter by measuring the cell capacitance by Patch-clamp. siM8-10 transfected cells were 1.3-fold bigger that other groups (**Fig. 3B**). We next confirmed the changes in GA morphology by immunolabeling both ERGIC and GA. No visible changes in GA morphology or surface area could be seen 2 days after siRNA transfection but expansion of ERGIC, cis-GA and TGN were obvious after 4 days (**Fig. 3C and Fig. S5A, S5B and S5C**). Quantitation of GA surface area confirmed a significant increase in siM8-10 group (**Fig. 3D**). The use of an alternative siRNA to siM8-10 confirmed the GA expansion while the use of a mutated siM8-10 prevented this effect (**Fig. S5B**). By contrast, no obvious changes could be observed in the ER, lysosomes or early endosomes (**Fig. S6**).

**FIGURE 3.**
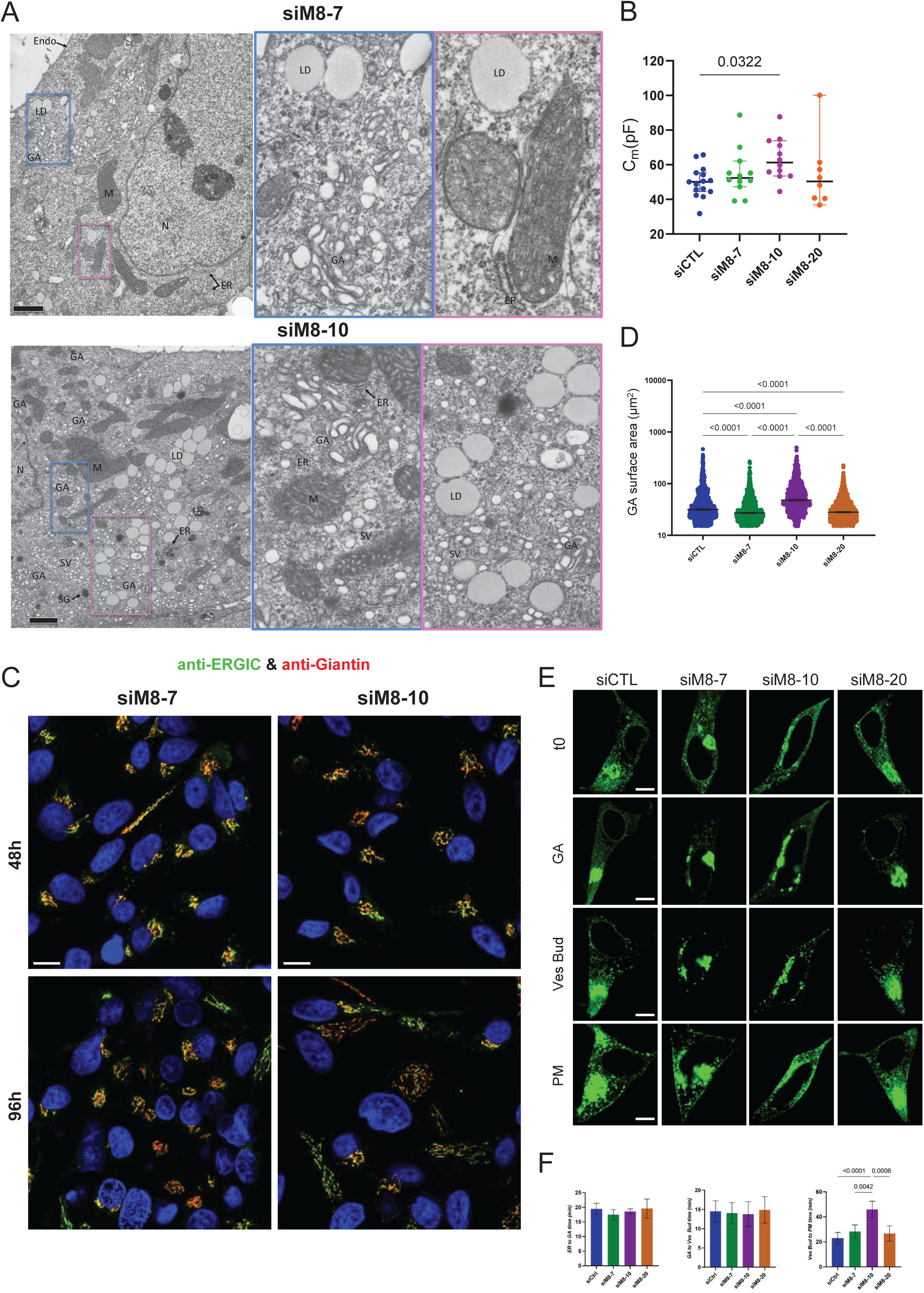
Induction of Golgi expansion and fragmentation in secretory vesicles and accumulation of lipid droplet in TRPM8(85) knock-down cells require expression of 4TM-TRPM8. **A,** TEM micrograph of cells subjected to group silencing of TRPM8 proteins. Labels: nucleus (N), endoplasmic reticulum tubules (ER), mitochondria (M), Golgi apparatus (GA), secretion vesicles (SV), secretory granules (SG), lipid droplet (LD), Black scale bars: 1 µm. **B,** Membrane capacitance values measured Patch-clamp recordings in siCTL, siM8-7, siM8-10, siM8-20 on LNCaP cells (96 hours after transfection) using the whole-cell configuration. **C,** Immunofluorescent labelling of Giantin (red) and ERGIC-53 (green) in LNCaP C4-2b cells transfected with siRNA for either 48h or 96h. Images were acquired with a confocal microscope. Scale bar: 10 µm. **D,** Quantitation of the GA surface area by after 96 hours of transfection. Cells were marked with Golgin 97 (TGN marker) antibody. Each point in the graph corresponds to surface area of one cell. Golgi surface area was measured using Image J. n=3 independent experiments. y-axis values are shown as log10 GA surface area and presented as Median ± 95% CI. Kruskal Wallis test was performed with p value< 0.05 considered significant. **E,** VSVG-GDP assay reports the effects of TRPM8 and TRPM8 isoform silencing on the protein trafficking from ER to plasmalemma in LNCaP cells. After 24 hours of siRNA transfection, cells were transfected with VSVG-GFP plasmid and then maintained in culture. Then after 24 hours of VSVG- GFP transfection, cells were switched to an incubator maintained at 40°C. After 12 hours, cells were shifted to 32°C and live cell imaging was recorded for minimum of 2 hours for each condition for tracking the VSVG-GFP transport. GA: Golgi apparatus; Ves Bud: beginning of vesicle budding; PM: Plasma membrane. Scale bar 10µm **F,** Time (min) of trafficking from ER to Golgi apparatus (GA), Golgi apparatus (GA) to vesicle budding (Ves Bud) and vesicle budding (Ves Bud) to plasmalemma (PM). Data are presented as Mean±SD. One-way ANOVA and Tukey’s multiple comparison tests were performed with p value< 0.05 considered significant. n=5 independent experiments were performed.

Secretory vesicles bud from the TGN and express some of TGN making it difficult to distinguish them from an expanded and fragmented GA. However, since secretory vesicles are transported from GA to plasmalemma, we hypothesized that an increase in their transport time could sign their accumulation in the cells. Using the vesicular stomatitis virus VSVG-GFP transport assay^48^, we quantified the transport times from ER to GA, GA to vesicle budding and vesicle budding to plasmalemma (**Fig. 3E)**. The co-silencing of TRPM8 and TRPM8(85) forms had an average transport time for vesicle budding to plasmalemma of 46±6 min while it ranged from 23±5 to 28±5 for the three other groups (**Fig. 3F**). We also assessed the effect of WS-12, a TRPM8 agonist, and BCTC and THIQj, two antagonists on VSVG-GFP transport times. Activation of TRPM8 channels decreased the time from GA entry to vesicle budding from about 15 to 10min. Inhibition of TRPM8s confirmed the increase in the average transport time for vesicle budding to plasmalemma of about 45min (**Fig. S7**).

In conclusion, these results showed that TRPM8 isoforms mitigated one or several mechanisms participating to GA structure, lipid droplets and vesicles exportation to plasmalemma. Common known features to these cell processes are lipid metabolism and membrane remodelling.

### TRPM8 isoforms shift the unsaturation of lipids, fatty acids chain length, and the choline to ethanolamine ratio

In order to identify the mechanism involved in the phenotype of siM8-10 transfected LNCaP cells and because the GA structure is finely-tuned by the lipid nature of its membranes^49^, we performed a high-throughput lipidomic screening by quantifying: cholesterol, free fatty acids (FFA), Phosphatidyl Choline (PC), Phosphatidyl Ethanolamine (PE), Phosphatidyl Inositol (PI), Phosphatidyl Serine (PS), Lysophosphatidyl Choline (LPC), Lysophosphatidyl Ethanolamine (LPE), Sphingomyelins (SM), Ceramids and dihydro- Ceramides (Cer and dCer) and plasmalogen Phosphatidyl Ethanolamine (pPE) in LNCaP C4-2b cells transfected with either siCTL, siM8-7, siM8-10 or siM8-20. The weight of each lipid family was normalized to the mean value of the siCTL group to highlight the changes induced by TRPM8s silencing. Cholesterol, total FFA, total PC, total PE, total LPC and total LPE, total SM and total pPE were all increased in the siM8-10 group (**Fig. 4A)**, as were total SFAs, total MUFAs and total Δ9-MUFAs, total PUFAs and long and very long chain FFAs. These results indicated that major changes occurred in LNCaP cells under the co-silencing of TRPM8 and TRPM8(85). The raw measurements of each individual lipids are reported in **Table 1**, and after normalization by cell size in **Table 2**. This normalization emphasized the difference in lipid content induced by shift in the lipid metabolism. For example, this revealed that very long chain ceramides were increased in siM8-10 independently of cell size (**Table 2**). Noteworthily, siM8-7 had a significant but mild effect on many lipid entities, which were further upregulated by siM8-10. This suggests that the shift in the lipid content depended on the silencing of both TRPM8 and TRPM8(85). Furthermore, suppression of all TRPM8 isoforms by siM8-20 reversed almost all the changes observed with siM8-10, suggesting that 4TM-TRPM8 expression was required for the phenotype observed in the siM8-10 group, consistent with the GA surface results. Beyond paired comparison of single lipid entities within groups, principal component analysis confirmed that siM8-10 group is the most different from the three others but it also showed that siM8-20 did not really recapitulate the control group (**Fig. 4B)**. Analysis of the main contributors of the first and second dimensions revealed that increase in Δ9 mono-unsaturation, poly- unsaturation and in PI contents are the main drivers of siM8-10 group while increase in SFA, PC and ceramides contents are the main contributors of siM8-20 group (**Fig. 4C)**. The third component of the PCA analysis segregated siM8-7 group (**Fig. S8A**) and one of the main contributors was the oleate (C18:1) incorporated in phosphatidyl groups in addition with ceramides (**Fig. S8B**).

**FIGURE 4.**
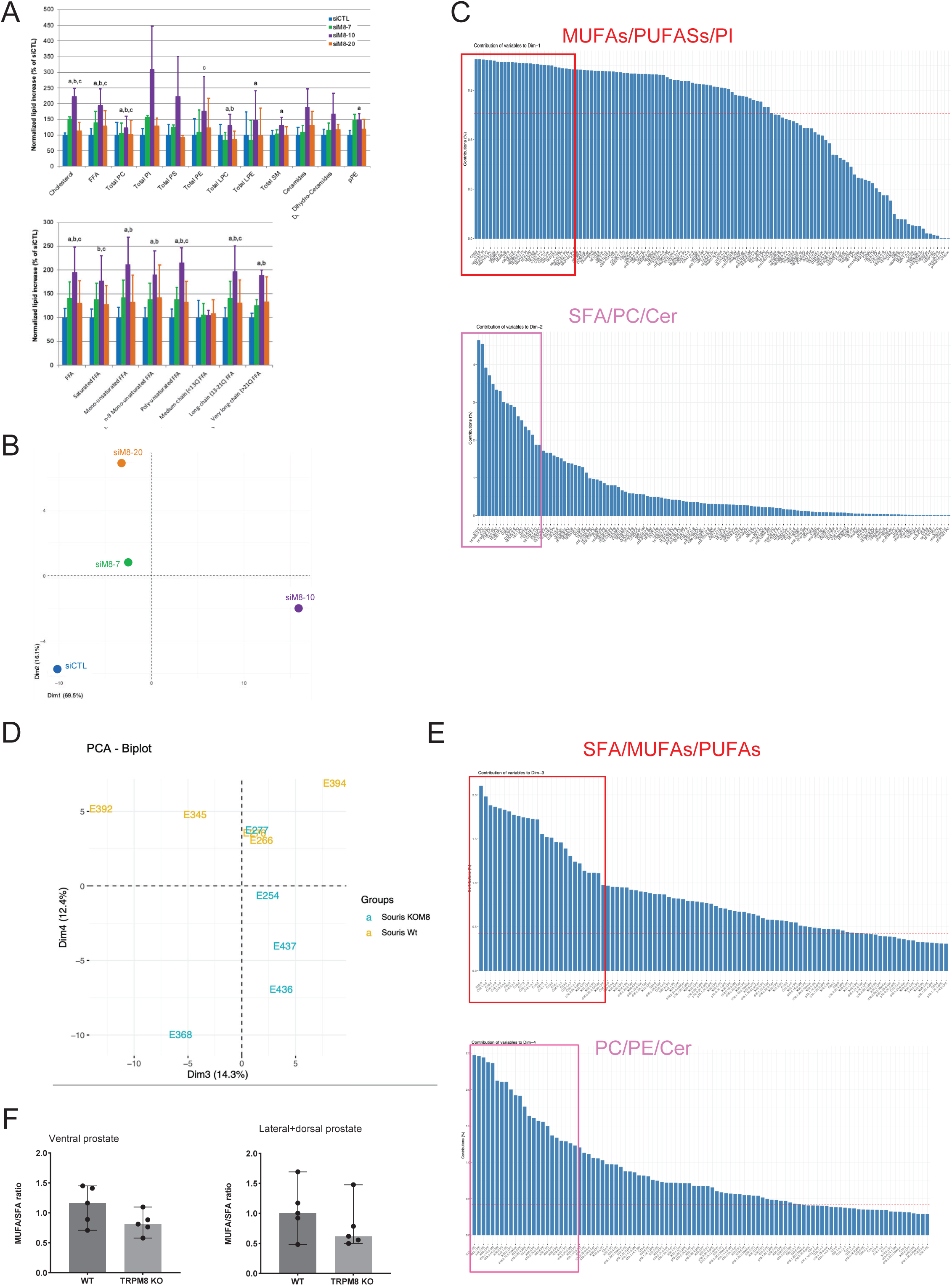
TRPM8 channels regulate unsaturation fatty acids, phospholipids and ceramides content in LNCaP C4-2b cells. **A,** Lipidomic analysis of LNCaP cell content after 96 hours transfection with siRNA targeting TRPM8 and its isoforms. Assessment of the content of different lipid species in LNCaP cells after siRNA treatment. Values of siM8 groups were normalized by the values of siCTL group (blue). (FFA: Free Fatty Acids, PC: Phosphatidylcholine, LPC: Lysophosphatidylcholine, PE: Phosphatidylethanolamine, LPE: Lysophosphatidylethanolamine, PI: Phosphatidylinositol, PS: Phosphatidylserine, SM: Sphingomyelin, pPE: Plasmenylethanolamine). The total level of saturation, unsaturation and chain length of FFA were also analysed (bottom graph). Values show: Mean±SD. Pairwise t-test was done to test siM8-10 group against siCTL (a: p<0.05), siM8-7 (b: p<0.05) and siM8-20 (c: p<0.05). **B,** Principal component analysis (PCA) biplot presents the divergency between siM8 groups scattered by the mean variance of their lipid species. Only the first and second dimensions (Dim1 and Dim2) are depicted since they explained 85% of the variance and split efficiently siM8-10 and siM8-20 groups. **C,** (top panel) Contribution of lipid species in dim1 (top panel) and dim2 (bottom panel). The red square and pink squares focused on the major contributors of each dimension in order to highlight the most represented lipids. The main contributors in dim1 and thus explaining the variance in siM8-10 group are: monounsaturated fatty acids (MUFAs), polyunsaturated fatty acids (PUFAs) and phosphatidylinositol (PI). The main contributors in dim2 and thus explaining the variance in siM8-20 group are: saturated fatty acids (SFA), phosphatidylcholine (PC) and ceramides (Cer). **D,** PCA biplot depicting the interplay between the lipid species and the mice (WT or KO M8 mice). Each of the points represents one mouse. Dimension 3 and 4 are presented because that enable the clearest scattering between *wt* and KOM8 mice. **E,** (top panel) Main lipid contributors to the dimension 3 of the PCA are shown in the red square: saturated fatty acids (SFA), monounsaturated fatty acids (MUFA) and polyunsaturated fatty acids (PUFA) and explained inter-individual variations between *wt* mice. (bottom panel) Main lipid contributors to the dimension 4 of the PCA are shown in the pink square: phosphatidylcholine (PC), phosphatidylethanolamine (PE) and ceramides (Cer) and explained the difference between *wt* and KOM8 mice. **F,** Saturation index: monounsaturated Fatty Acids (MUFAs) to saturated fatty Acids (SFAs) ratio in ventral and lateral/dorsal prostate from WT and TRPM8 KO mice. Pairwise comparison with t-test has been done (p=0.11 and 0.18, in ventral and latero-dorsal prostates, respectively) Data are presented as Median±95%CI.

**Table 1.**
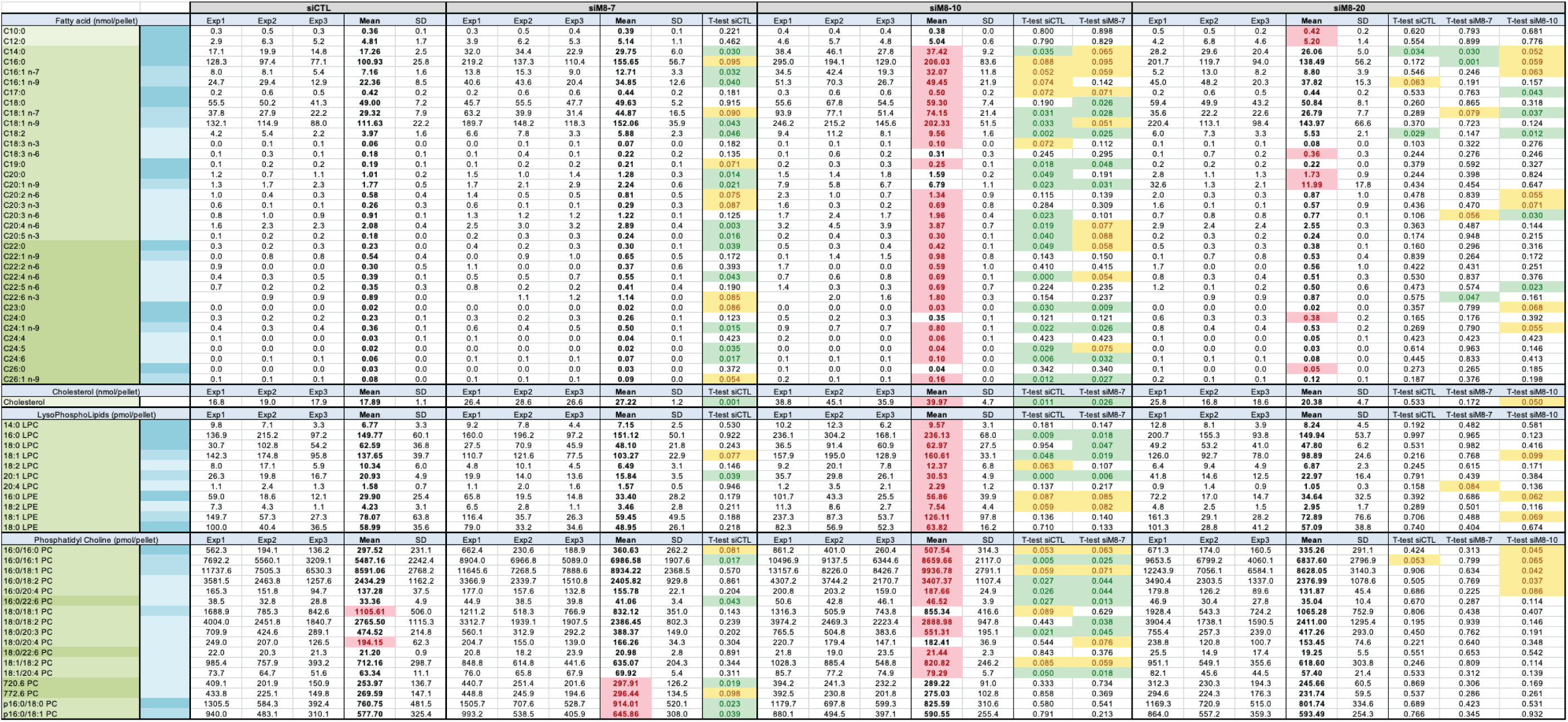
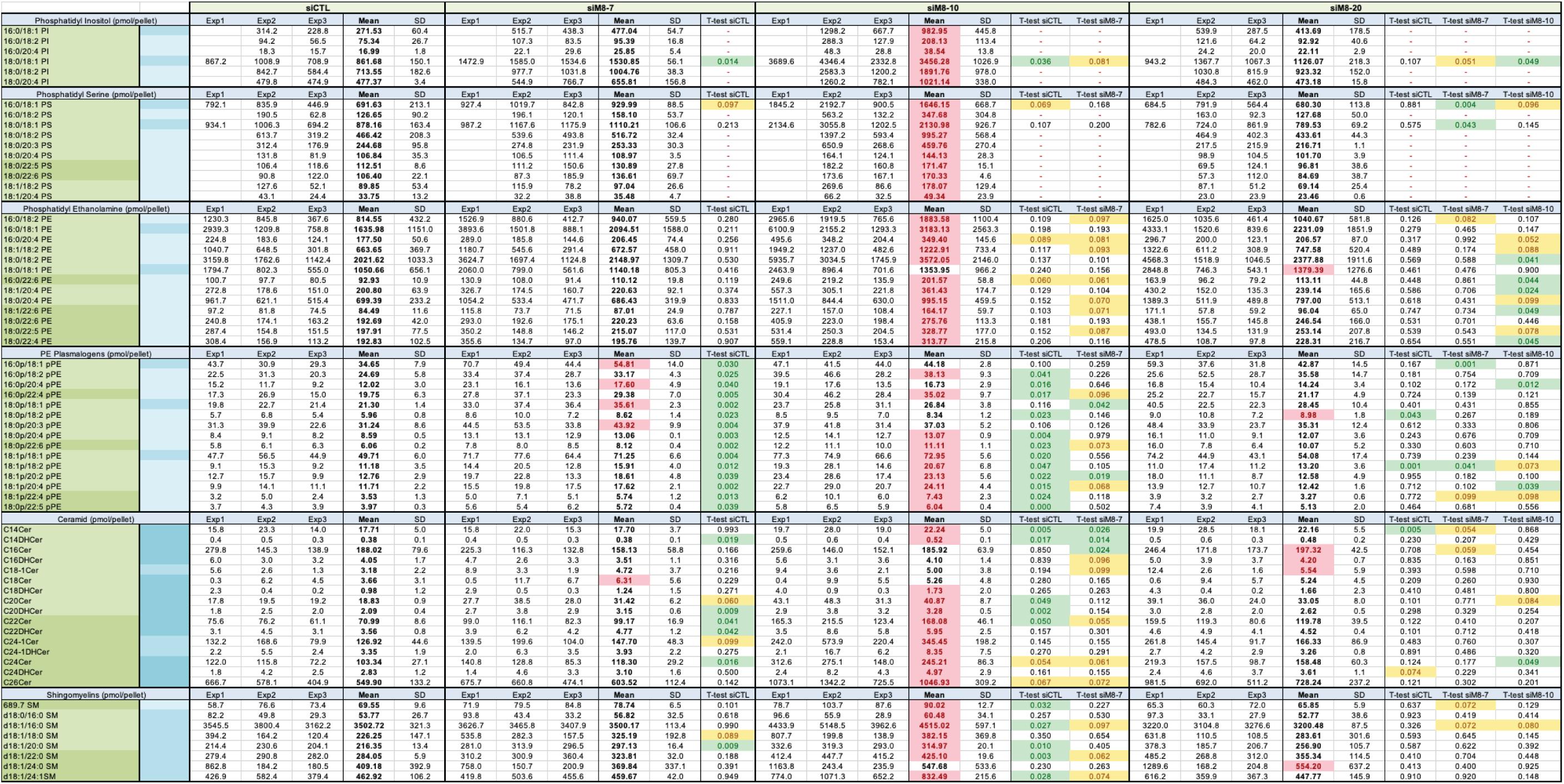
is reporting the raw values of lipid content per cell pellets (2 million cells per pellet) for each class of lipids in the four groups (siCTL, siM8-7, siM8-10, siM8-20) of LNCaP C4-2b cells. The first column shows the different lipids analyzed and sorted by class. The color code in the first column reports the fatty acid chain length: medium (<C14), long (C14-C21) and very long (>C21), from light and medium green to dark green, respectively. The color code in the first column reports the saturation level of the acid chain (sn2 position for CER and SM): saturated, mono-unsaturated and poly-unsaturated, from dark and medium blue to light blue, respectively. Values from 3 independent experiments are shown, excepting several measures for PI and PS assays that have been excluded because of the detection of contaminant in the experimental control. Mean and standard deviation (SD) are shown and the highest mean value amongst the four groups is highlighted with red fond and red cell background. T-test against the indicated groups are showed on the right of each group and significant test (below 0.05) are highlighted with green font and green cell background. Pairwise comparison is lacking sensitivity because of the limited statistical power. Therefore, p values between 0.05 and 0.1 are given highlighted with yellow font and yellow cell background for information purposes.

**Table 2.**
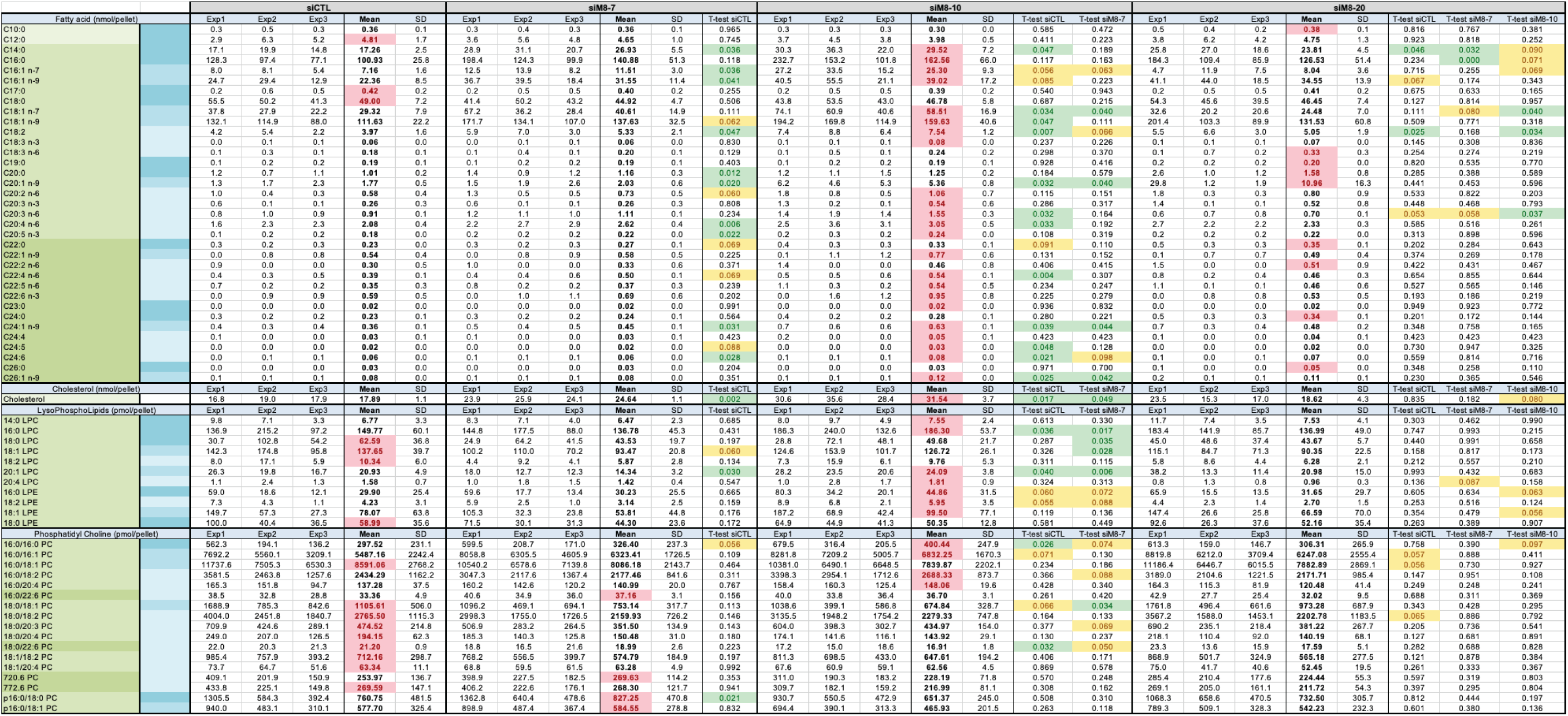
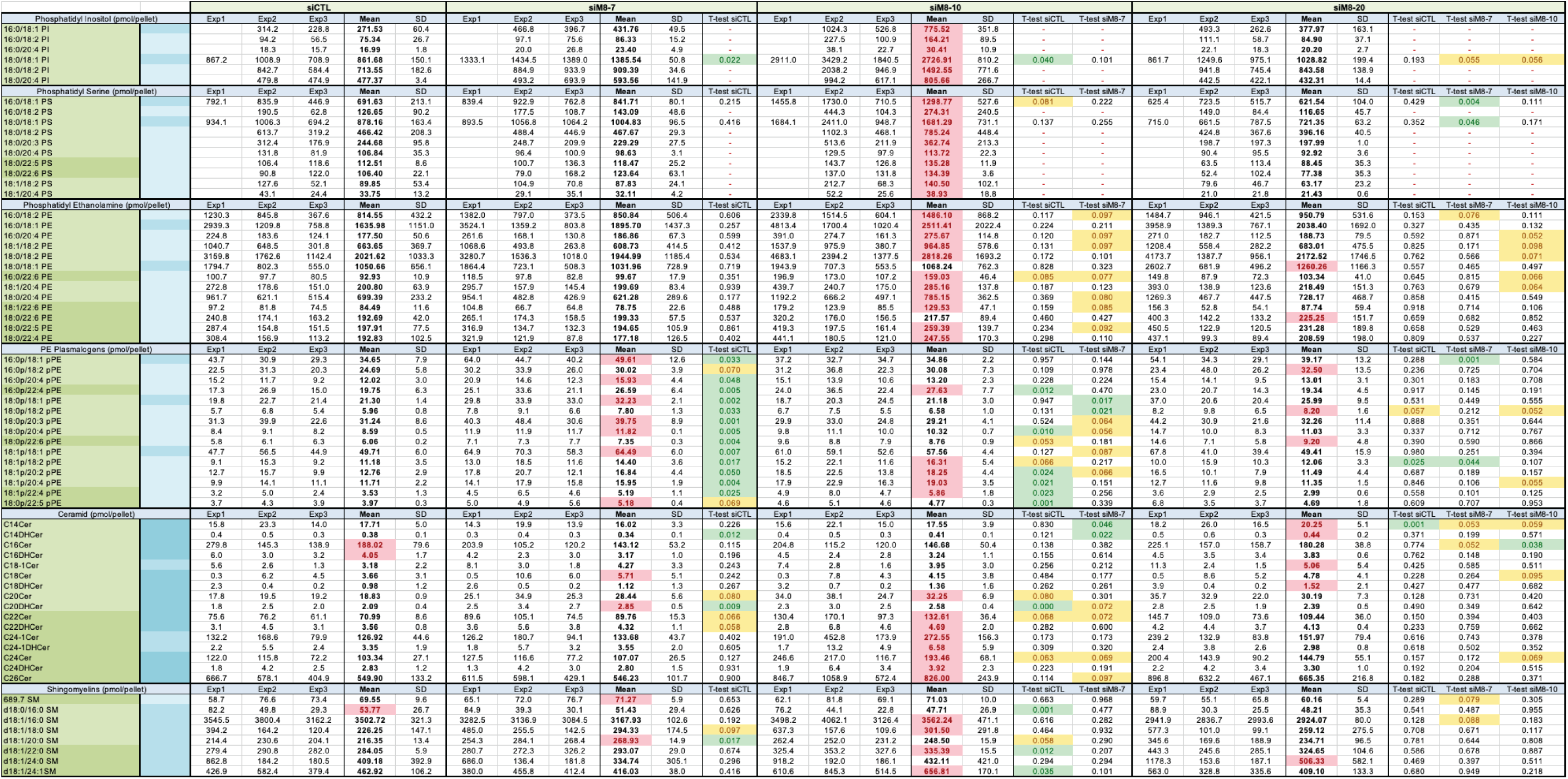
is reporting the normalized values of lipid content per cell size for each class of lipids in the four groups (siCTL, siM8-7, siM8-10, siM8-20) of LNCaP C4-2b cells. The first column shows the different lipids analyzed and sorted by class. The color code in the first column reports the fatty acid chain length: medium (<C14), long (C14-C21) and very long (>C21), from light and medium green to dark green, respectively. The color code in the first column reports the saturation level of the acid chain (sn2 position for CER and SM): saturated, mono-unsaturated and poly-unsaturated, from dark and medium blue to light blue, respectively. Values from 3 independent experiments are shown, excepting several measures for PI and PS assays that have been excluded because of the detection of contaminant in the experimental control. Mean and standard deviation (SD) are shown and the highest mean value amongst the four groups is highlighted with red fond and red cell background. T-test against the indicated groups are showed on the right of each group and significant test (below 0.05) are highlighted with green font and green cell background. Pairwise comparison is lacking sensitivity because of the limited statistical power. Therefore, p values between 0.05 and 0.1 are highlighted with yellow font and yellow cell background for information purposes.

Since siM8-20 inhibited all forms of TRPM8, we expected its phenotype to be similar to the phenotype of prostate epithelial cells in KOM8 mice. PCA confirmed the segregation of *wt* and KOM8 samples mainly along the fourth component while the third component segregated samples inside the *wt* group (**Fig. 4D**). The segregation between the *wt* and KOM8 groups was mainly attributable to changes in PC and PE species. Variation in SFA, MUFA and PUFAs contents were mostly related to differences between *wt* mice (**Fig. 4E**). Even if not significant because of a lack of power, the pairwise comparison between MUFA/SFA ratio reported a downward trend (**Fig. 4F**). Values for each lipidic entity, along with the mean±SD and pairwise t-test are reported in **table 3 and 4** for ventral and latero-dorsal prostates, respectively.

**Table 3.**
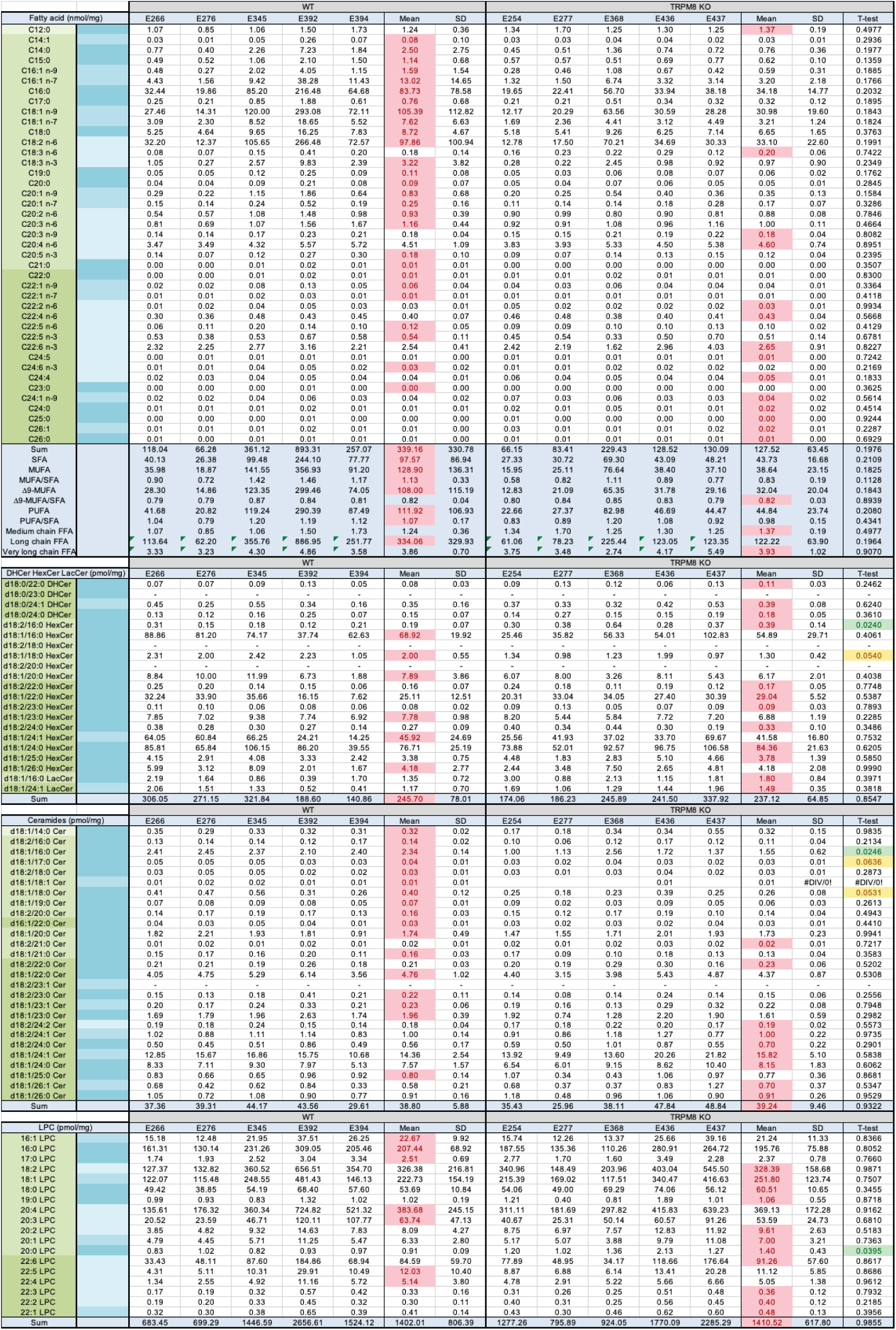
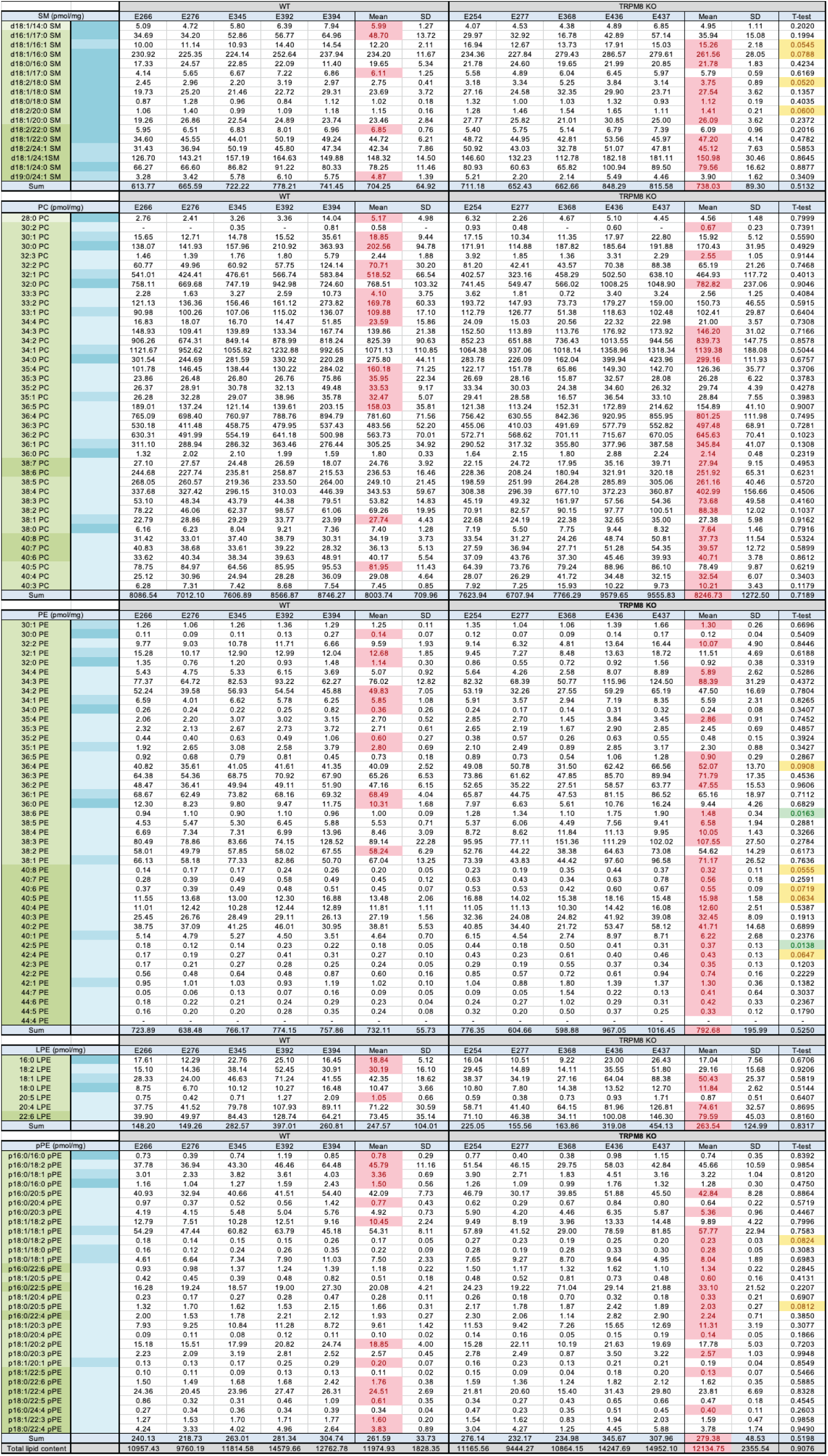
is reporting the raw values of lipid content mg of ventral prostate from either *wt* (littermates) or KOM8 mice. The first column shows the different lipids analyzed and sorted by class. The color code in the first column reports the fatty acid chain length: medium (<C14), long (C14-C21) and very long (>C21), from light and medium green to dark green, respectively. The color code in the first column reports the saturation level of the acid chain (sn2 position for CER and SM): saturated, mono-unsaturated and poly-unsaturated, from dark and medium blue to light blue, respectively. Values from 3 independent experiments are shown, excepting several measures for PI and PS assays that have been excluded because of the detection of contaminant in the experimental control. Mean and standard deviation (SD) are shown and the highest mean value amongst the four groups is highlighted with red fond and red cell background. T-test against the indicated groups are showed on the right of each group and significant test (below 0.05) are highlighted with green font and green cell background. Pairwise comparison are lacking sensitivity because of the limited statistical power. Therefore, p values between 0.05 and 0.1 are given highlighted with yellow font and yellow cell background for information purposes.

**Table 4.**
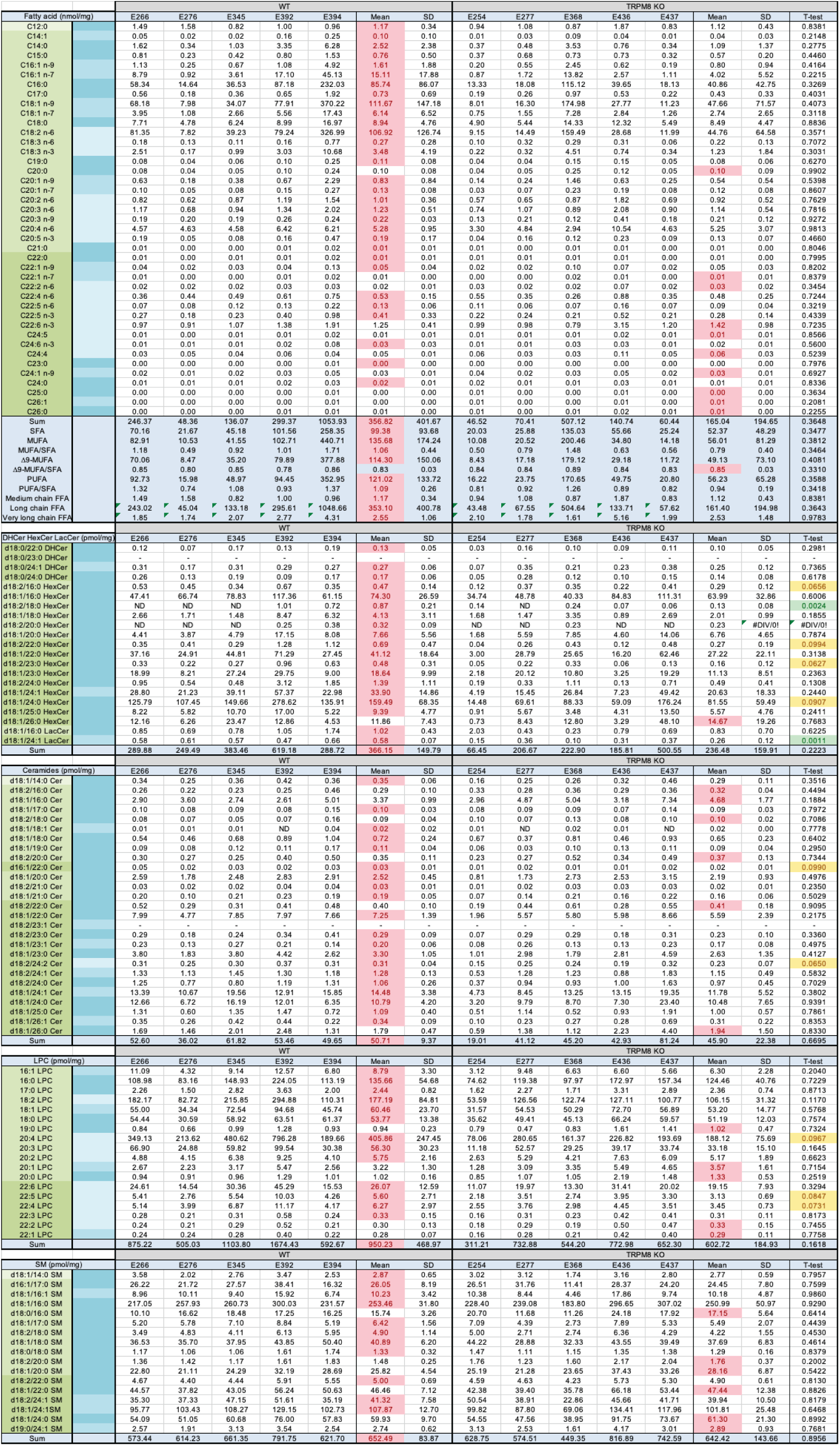
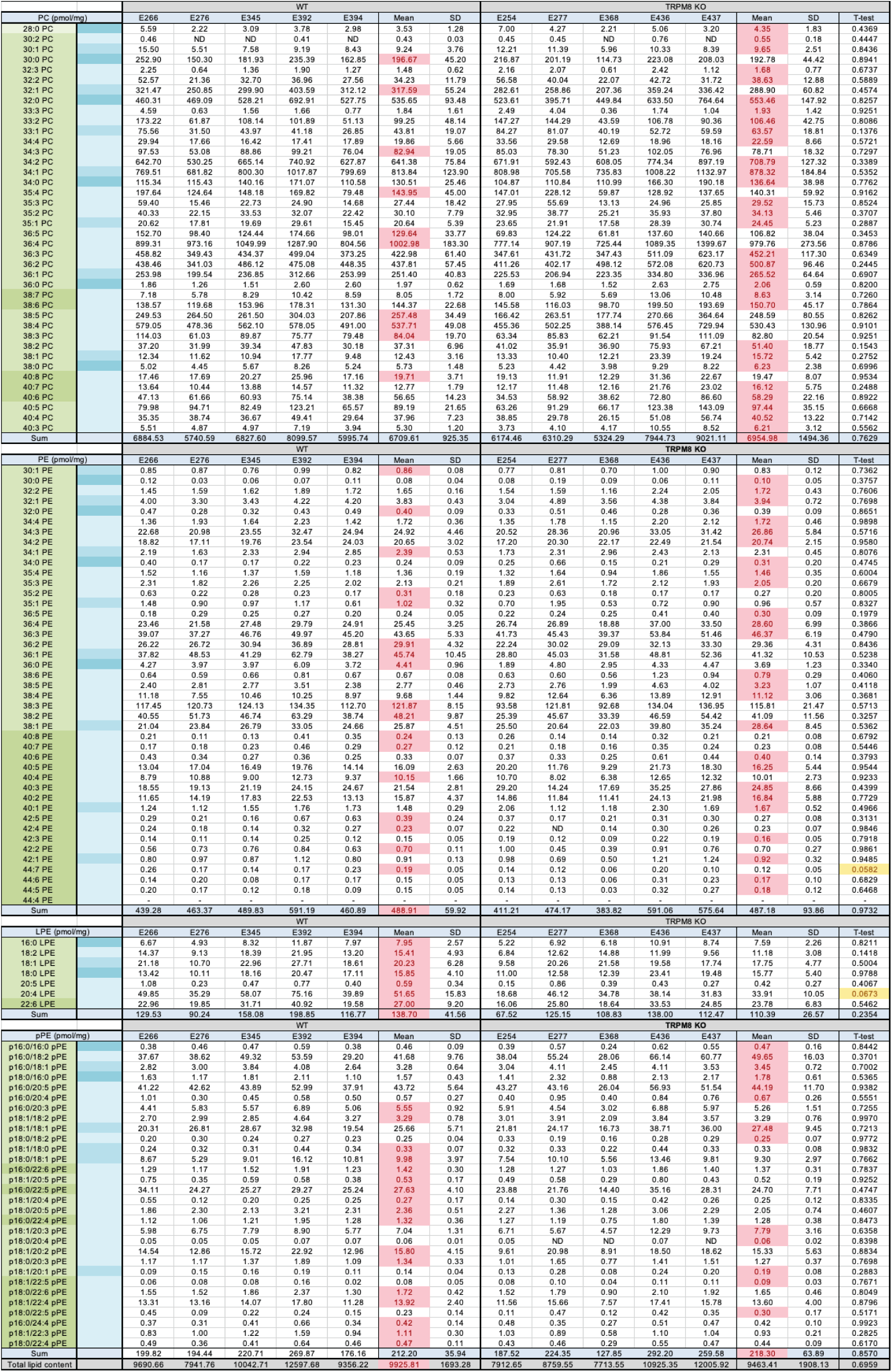
is reporting the raw values of lipid content mg of dorsal and lateral prostate from either *wt* (littermates) or KOM8 mice. The first column shows the different lipids analyzed and sorted by class. The color code in the first column reports the fatty acid chain length: medium (<C14), long (C14-C21) and very long (>C21), from light and medium green to dark green, respectively. The color code in the first column reports the saturation level of the acid chain (sn2 position for CER and SM): saturated, mono-unsaturated and poly-unsaturated, from dark and medium blue to light blue, respectively. Values from 3 independent experiments are shown, excepting several measures for PI and PS assays that have been excluded because of the detection of contaminant in the experimental control. Mean and standard deviation (SD) are shown and the highest mean value amongst the four groups is highlighted with red fond and red cell background. T-test against the indicated groups are showed on the right of each group and significant test (below 0.05) are highlighted with green font and green cell background. Pairwise comparison are lacking sensitivity because of the limited statistical power. Therefore, p values between 0.05 and 0.1 are given highlighted with yellow font and yellow cell background for information purposes.

Altogether, these results reported that 1) TRPM8(85) silencing increased ceramides level with very long chain FA at the sn2 position, 2) suppression of all TRPM8 isoforms did not recapitulate the control prostate but showed a decrease in PC content, compensated by an increase in PE content, 3) suppression of TRPM8 induces an increase in FFA unsaturation and in oleate at sn1 position of SM, which was drastically enhanced by the additional deletion of TRPM8(85).

### TRPM8 isoforms control SCD1 expression in an AR-dependent mechanism

Since i) the co-silencing of TRPM8 and TRPM8(85) had major effects on the unsaturation level of FFAs, ii) MUFA/SFA ratio is the key component of the cold sensing feature of DesK system in bacteria, iii) PUFAs have been shown to inactivate TRPM8^36^, we decided to continue investigating the role of TRPM8 on the rate-limiting enzyme of MUFA generation: the Δ9 stearoyl desaturase 1, SCD1. Immunoblotting showed an increase in SCD1 expression in siM8-10 group (**Fig. 5A and 5B**). We additionally assessed the Fatty Acid Synthase (FAS) which is reported to usually work in tandem with SCD1 and we found that its expression was unchanged under siM8s transfection (**Fig. S9A and S9B**). This suggested to some extent that TRPM8 isoforms exerted a specific regulation on lipid remodelling rather than not on lipid synthesis. SCD1 has been proposed to be located in the ER membranes since its enzyme activity was measured in microsomal preparations^50^. However, microsomes include not only ER membranes but also vesicles of similar density. In addition, to explain why siM8-10 induced GA expansion, we cloned human SCD1 and fused it in frame with the YFP prior to co-expressing it with a SERCA-mT2 protein in LNCaP C4-2b cells. As shown, in figure 5C, although fluorescence of SCD1 can be partially colocalized with SERCA, it was much concentrated in one restricted area around the nucleus as expected of the GA (**Fig. 5C**). We found that SCD1 silencing could prevent GA expansion in siM8-10 transfected cells (**Fig. 5D**) but that silencing SCD1 alone did not change the GA surface area. To the opposite, overexpression of SCD1-YFP increased the GA surface area independently of the expression of the canonical TRPM8 (**Fig. 5E**). Because modulation of SCD1 expression is expected to trigger changes in MUFA/SFA ratio, we hypothesized that changes in MUFA/SFA ratio is the driving mechanism for GA expansion. We incubated transfected LNCaP C4-2b cells with either palmitate or oleate. Palmitate incubation was able to revert siM8-10-induced GA expansion which confirmed that a relative decrease in SFA is required to induce an increase in GA surface area (**Fig. 5F**). Oleate had not additional effect on GA surface area in siM8-10 group but it did induce GA expansion in control cells (**Fig. 5F**). Surprisingly, oleate decreased GA surface area in siM8-7 group. These results confirmed that the increase MUFA/SFA ratio was the driver of GA expansion.

**FIGURE 5.**
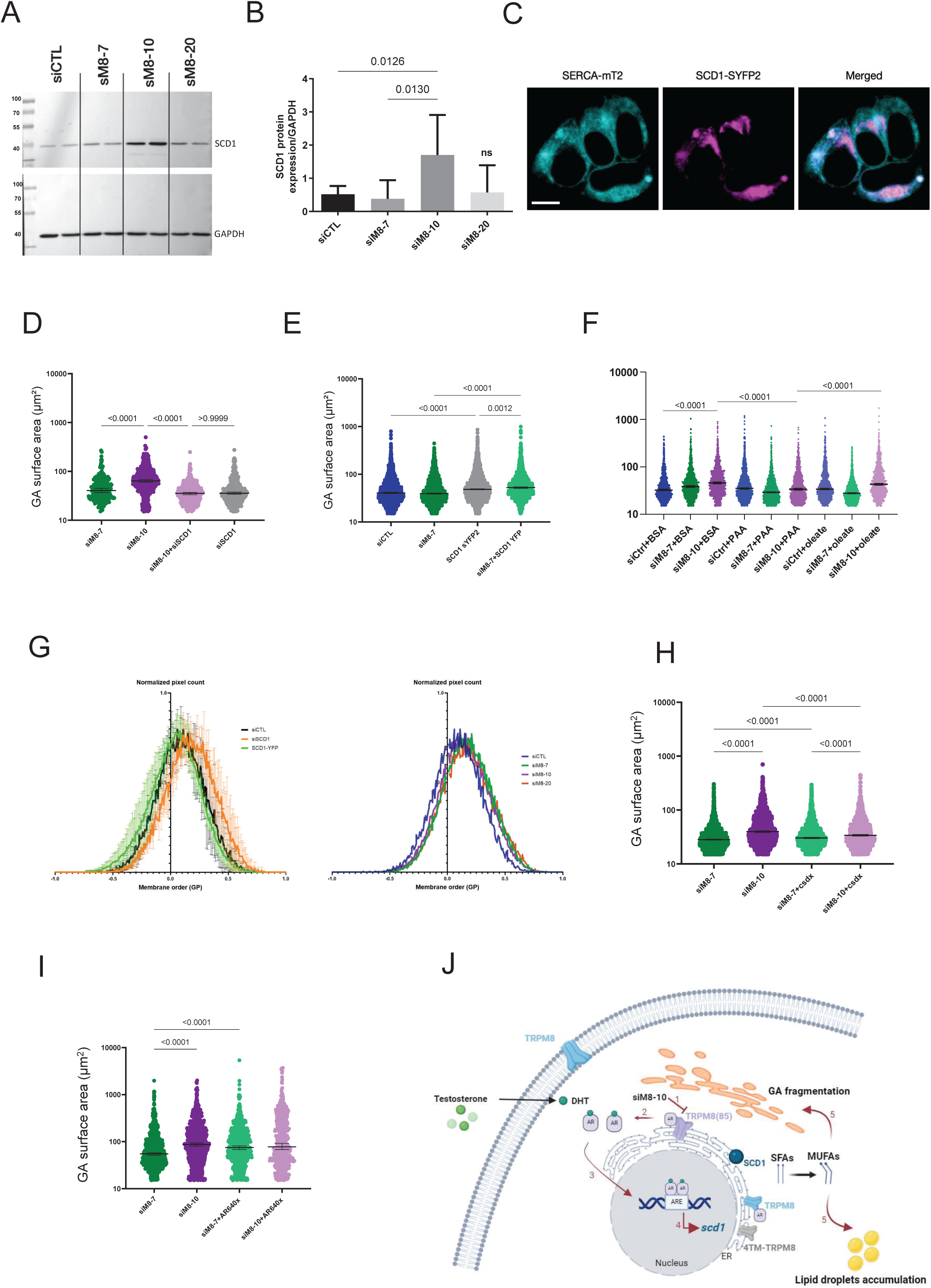
TRPM8 silencing upregulates SCD1 expression by an AR-dependent mechanism, increases the MUFA/SFA ratio that triggers GA expansion. **A-B,** Western-blot shows an increased SCD1 protein expression in LNCaP C4-2b cells transfected with siM8-10 for 2 days. After 48 hours of siRNA transfection, SCD1 protein expression was assessed by western blot. n=5 independent experiments. Median±95%CI. Kruskal Wallis test was performed with the indicated p-values. **C,** Confocal images of LNCaP cells expressing SERCA-mTurquoise2 (blue ER marker) and SCD1-SYFP2 (pink) proteins. Overlay of the two fluorescent signals is shown in the 3^rd^ column. Scale bar 10µm. **D, E, an F,** Quantitation of GA surface area by after 96 hours of transfection. LNCaP cells were co-transfected with siM8-7 or siM8-10 along with siSCD1 (**D**), co-transfected with siCTL and siM8-7 along with SCD1 (**E**) or LNCaP cells were transfected with siCTL, siM8-7 or siM8-10 and incubated with BSA, 50µM palmitate or 50µM oleate (**F**). Cells were labelled with anti-Golgin-97 (TGN marker) antibody. Each point in the graph corresponds to surface area of one cell. Golgi surface area was measured using Image J. **G,** Membrane order analysis by the generalized polarisation method using Laurdan (6-Dodecanoyl-2-dimethylaminonaphthalene), a lipophilic dye capable of partitioning into cell phospholipidmembranes. (left panel) LNCaP cells were transfected with siCTL, siSCD1 or SCD1-YFP plasmid for 48 hours. (right panel) LNCaP cells were transfected with siM8-7, siM8-10, or siM8-20 for 48 hours. **H and I,** Quantitation of the Golgi surface as in panel D for LNCaP cells were transfected with siM8-7 or siM8-10 and either treated with androgen receptor blocker, bicalutamide (casodex®; csdx) at 10µM for 72 hours (**H**) or co-transfected with an AR-positive mutant, AR640X (**I**). In the panels assessing GA surface area (D,E,F,H and I), data are presented as log_10_ Golgi surface area on the y- axis. n=3 independent experiments. Median±95%CI. Kruskal Wallis test with p value< 0.05 considered significant. **J,** Schematic representation of the newly characterized TRPM8(85) isoform in the ER and its involvement in the control of GA morphology and activity through the fine tuning of GA Ca^2+^ content and the saturation/desaturation ratio of fatty acids. TRPM8(85) is localized in the ER at the interface with the GA. Knocking down TRPM8(85) by siM8-10 leads to AR dimer translocation into the nucleus which then binds androgen responsive element (ARE) in the promoter region of the *Scd1* gene activating its transcription. SCD1 protein, residing in ER and GA, then catalyses the reaction of MUFA synthesis: palmitoleate and oleate, from SFA: palmitate and stearate, respectively. This increases the level of MUFA in GA induces its fragmentation and expansion due modification in membrane biophysical properties other than membrane order. Additionally, the increased MUFA or decrease in PC/PE ratio in the cells could be responsible for lipid droplets formation and accumulation in the cells.

An increase in the level of membrane unsaturation is known to increase membrane order^51^, which has been proposed to be the main mechanism controlling cold sensing of the DesK system in bacteria^27^. By analysing the generalized polarisation (GP) of the fluorescent Laurdan probe^38^, we assessed membrane order in LNCaP C4-2 cells. First, we were unable to measure variations in GP in cells incubated with either Palmitate or Oleate (**Fig. S11**), suggesting that this treatment was not sufficient to modify the average membrane order in the cells. Nevertheless, SCD1-silenced cells displayed a strong increase in GP values distribution, conversely SCD1-YFP expression slightly decrease membrane order (**Fig. 5G**). These results validated that focused modification on SCD1 over-expression correlated with the expected change in membrane order and confirmed that a decrease in membrane order is sufficient to induce GA expansion (**Fig. 5D and 5E)**. Intriguingly, siM8-treated cells showed a similar GP feature showing a slight increase in membrane order as compared to control cells. However, no clear difference in membrane order was observed between siM8-10 and the two other siM8 groups (**Fig. 5G**). In conclusion, we believe that changes in membrane order did not account for GA expansion induced by the co-silencing of TRPM8 and TRPM8(85).

We showed that SCD1 is a key enzyme involved in the effect induced by co-silencing of TRPM8 and TRPM8(85) and that modification in the SFA/MUFA ratio is involved. We finally wondered by which mechanism TRPM8 could regulate SCD1 expression. In 2018, Audet-Walsh et al. reported that AR and mTor could induce the expression of the sterol regulatory element-binding transcription factor 1 which in turn induces FAS and SCD1 expression^52^. Furthermore, TRPM8 has been demonstrated to be a non-genomic target of AR^53^ whose binding inhibited TRPM8 channel activity^54^. We thus assessed if AR could be involved in TRPM8-mediated regulation of SCD1 expression. AR inhibition by bicalutamide inhibited siM8-10- mediated GA expansion (**Fig. 5H**) but had no effect on siM8-7 group. Transfection of the constitutive nuclear AR mutant (AR640X) induced a significant GA expansion in siM8-7 transfected cells but had no additional effect on siM8-10 group (**Fig. 5I**).

Altogether, our results reported a complex regulation of lipid metabolism by TRPM8 and its isoforms. We especially demonstrated the role of TRPM8 and its isoform in the physiological control of GA morphology through the fine tuning of SCD1 expression and the MUFA/SFA balance (**Fig. 5J**).

## DISCUSSION

### Calcium regulation and TGN export rate to plasmalemma

SPCA1-mediated Ca^2+^ uptake in GA has been shown to drive the transport of proteins within GA compartments and their final export out of GA^55^. We observed an increase in the steady-state Ca^2+^ concentration in the TGN in TRPM8(85)-silenced cells that cannot be explained by either TRPM8- or TRPM8(85)-dependent increase in ER-to-TGN Ca^2+^ transfer. However, we cannot exclude the possibility that 4TM-TRPM8 could be involved in transferring Ca^2+^ from ER-to-Golgi. Indeed, we reported in the current study that all the effects induced by the co-silencing of TRPM8 and TRPM8(85) required the expression of 4TM-TRPM8. Increase in GA Ca^2+^ content has been shown to trigger 1) the Ca^2+^-dependent sorting through Cab45 oligomerization^56^, and 2) activation of the calneuron 1 and 2 (also known as CBP8 and CaBP7) Ca^2+^-sensors^57^, what lead to the inhibition of phosphatidylinositol 4-kinases IIIB (PI4K IIIB) in TGN. Consequently, PI accumulates (as measured in our study) and is paralleled by a decrease in phosphatidyl Inositol-4-phosphate (PI4P) pools and a decrease in vesicle trafficking to plasmalemma (as measured in our study). We thus infer that TRPM8(85) and 4TM-TRPM8 isoforms are involved in the GA Ca^2+^ homeostasis of prostate epithelial cells that could participate in the orchestration of GA morphology and activity of vesicle secretion.

### Golgi apparatus morphology, the TRPM8-SCD1 connection

GA morphology has been observed since decades by electron microscopes and variety of structural TGN have been reported^58^. Although these studies have focused on static geometrical features, GA activity is highly dynamic requiring membrane synthesis and remodelling to compensate for the intense vesicle budding^59^. This constant remodelling of GA membranes is starting with, first, the synthesis and export of PC, PE, PI, Lysophospholipids, ceramides, SM and Cholesterol synthesis in ER^60^. Second, the distribution of enzymes, resident proteins and lipid composition in GA differs throughout, from the cis-Golgi to the trans-Golgi. PI are specifically concentrated in GA where they are used as precursor of PI4P. A decrease in PI4P in GA has been shown to down-regulate ER-to-GA transport of ceramides and the consecutive de novo synthesis of SM in GA. Both ceramides and SM were up-regulated in TRPM8(85)-silenced cells, which suggests that PI4P content was unchanged. Last component of the GA dynamics, vesicles budding from GA requires both specialized proteins and lipid modulating the fusion/fission processes orchestrated by proteins and lipids^61^. In our model, up-regulation of MUFA, PI, long chain ceramides and SM level was observed when silencing TRPM8(85) and to a lesser extent TRPM8 and could thus participate in a complex lipid-mediated induction of GA expansion and activity. However, it is experimentally difficult if not impossible, to assess each lipidic class to determine whether it is a cause or a consequence of GA expansion, and in what extent. We could achieve it for MUFA pathway since SCD1 is the rate-limiting enzyme.

The effects of an unbalanced MUFA/SFA ratio on GA homeostasis have been largely ignored in the literature. Lita et al reported that a mutation of the cytosolic isocitrate dehydrogenase 1 increased SFA-, and MUFA- PEs and PCs in ER but depleted them in GA. This depletion in GA was actually associated with an upregulation of SCD^62^. Differently from their results, we neither observed a dilation of ER nor of GA but rather an expansion and fragmentation of GA. In addition, we did not measure any modification in FAS expression level. Payet et al, reported that SFA (mainly C14:0, C16:0 and C18:0) accumulations in a mutated yeast model was associated with reduced vesicle formation in the TGN^63^. Along the opposite line, we found that accumulation of MUFAs and PUFAs was accompanied with a normal formation of vesicles but with a slower trafficking. Finally, we showed that silencing SCD1or incubation of cells with palmitate suppressed GA expansion, confirming the major role of MUFA/PUFA increase in the regulation of GA morphology.

### Beyond membrane order in TRPM8(85)-induced GA expansion

In the DesK cold sensing model in bacteria, the cold effect is mimicked by SFAs through their ability to compact the membrane, also known as an increase in membrane order. We observed an interplay between membrane order with a modification in GA surface area when overexpressing SCD1 and when incubating cells with oleate. Increase in MUFA is thus correlated with a decreased membrane order and with GA expansion. However, in TRPM8(85)-silenced cells, SCD1 was induced, MUFAs were increased and GA expanded but membrane order was not significantly changed - suggesting that membrane order is not involved. Numerous lipid species were modified in TRPM8(85)-silenced cells: MUFAs, PUFAs, Cer, SM, PC, PE, PS, PE or plasmalogen PE. Interestingly, plasmalogen PE have been reported to form thicker, compressed and rigid bilayers than their equivalent PE^64^ and thus could have compensated for MUFAs effect on membrane order. Beyond membrane order, it is known that the variation of several lipids like MUFAs, PC and PE modify membrane curvature and stability of GA membranes^65^. We thus cannot exclude that modification in lipids triggers a variation in membrane curvature that facilitates directly GA expansion and fragmentation.

### TRPM8s: beyond cold, a lipid sensor orchestrating membrane remodelling?

Beyond cold, TRPM8 has been shown to be activated by lysophospholipids and PI(4,5)P while being inhibited by PUFAs and cup-forming cationic amphipaths (PMID: 17082190; PMID: 17376995; PMID: 15852009; PMID: 32101066). Interestingly, LPC, which is known to decrease membrane order in PC membranes has been shown to potentiate cold activation of TRPM8^36^. Beyond its role as a cold sensor in tissues or cells exposed to environmental variation of temperature, we believe that TRPM8 and its isoforms being expressed in organs insulated by the body, should be considered as a lipid sensor able to regulate SCD1 expression and MUFAs content in prostate cancer cells.

### TRPM8-AR interplay in prostate cells

In 2004 and 2005, Barrit’s team and Prevarskaya’s team demonstrated the *Trpm8* is an AR-regulated gene^14,17^. Later, Zakharian’s team advocated that the TRPM8 channel function as a Testosterone-gated channel^66,67^ in absence of AR. Grolez et al demonstrated that PolyQ domain of AR interacts with both the amino-terminal and the carboxy-terminal loops of TRPM8 and is required for TRPM8 activity^53^. Although, the effect of AR-binding to TRPM8 gating has been studied in the context of activation by TRPM8 agonists, it is unknown whether activation by lipids (PIP2, Lysophospholipids) is also inhibited by AR-binding. Gradual increase in testosterone concentration weakens AR binding to TRPM8, leading to AR release from TRPM8 carboxy-terminal extremity which inhibits TRPM8^53^. Free AR is expected to be able to translocate into the nucleus and regulate its target genes such as *Trpm8*. In the current study, we reported that AR is involved in the transmission of information between TRPM8(85) and *Scd1* gene. In this TRPM8-AR model, decreased expression in TRPM8 proteins is equivalent to a decrease in binding traps for AR that should free it even in absence of testosterone and let it trans-activates *Scd1* gene in the androgen-resistant LNCaP C4-2b line. Conversely, the inhibition or decreased expression of AR should inhibit TRPM8 channels and disrupt the signalling towards *Scd1* gene. Along this line, a recent spatial-transcriptomic study revealed that anti-androgen responsive prostate cells of a patient showed a low TRPM8 expression along with a high SCD1 expression before the treatment^68^. Noteworthily, in castration-resistant cells TRPM8 was upregulated while SCD1 was down-regulated. We believe these independent results confirm our interpretation that TRPM8 expression levels determine a trapping force on AR that, in addition to testosterone levels or activating mutations on AR, determines the transcriptional activity of the AR-dependent genes like *Scd1*.

## Conclusion

In the current study, we revealed a new and original role for TRPM8 and its isoforms as modulators of SCD1 expression. This finding establishes their role in controlling GA membrane content and morphology, along with regulating the secretory vesicle transport.

## AUTHOR CONTRIBUTIONS

Conceived and performed experiments: MW, YG, FG, ASB, JG, SB, CC, CS, DG, ED, PD, JPPDB, LL, GB Wrote the manuscript: GB, YG, MW

Supervised the study: GB, YG, ASB

Provided expertise and feedback on FLIM experiments: LH Review the manuscript: RF, DG, MK, FVC, LL, NP

## AKNOWLEGDMENTS

This work was supported by grants from INSERM, Ministère de l’Education Nationale, Ligue Nationale Contre le Cancer, the Region Nord-Pas-de-Calais, SIRIC OncoLILLE, Association de Recherche sur les Tumeurs de la Prostate, IHU OPeRa (ANR-10-IBHU-004) program “Investissements d’Avenir” operated by the French National Research Agency (ANR), Leducq Transatlantic Network of Excellence “Targeting Mitochondria to Treat Heart Disease ‘MitoCardia” (16 CVD 04), INSERM and USTL annual funding of the PhyCell Laboratory, INSERM and UCBL annual funding of the CarMeN Laboratory.

Dr. Anne-sophie Borowiec was supported by Fondation pour la recherche médicale.

D1ER cameleon and 4mtD3cpv Cameleon are generous gift of Prof Tsien R., La Jolla, CA, USA. GoD1cpv cameleon is a gift from Dr Foulquier F (Univ. Lille, CNRS, UMR 8576 - UGSF - Unité de Glycobiologie Structurale et Fonctionnelle, F-59000 Lille, France).

Authors thank Dr Rialland M (INSERM UMR, 1231, Dijon, France) for his critical advices along the study and technicians of the DiviOmics platform (UMS 58 BioSanD, Université de Bourgogne - 21000 Dijon - France) for the lipidomic analysis.

## Supplementary information

## SUPPLEMENTAL DISCUSSION

### TRPM8(85): a primate story?

We previously reported the diversity of alternate TRPM8 mRNA^1^ and characterized several of these isoforms^2–5^. In addition to their characterization, we looked at the evolution of human TRPM8 forms and found that most of the alternate *Trpm8* exons were selected lately in evolution – showing conservation of splicing features in primates and a higher percentage of transposable elements in alternate exons than in core exons^1^. Comparatively, the core sequence of the *Trpm8* gene, coding for the canonical TRPM8 channel has been conserved since Terapoda^6^. These results did not preclude possible paralogous evolution in rodents, although some human alternate *Trpm8* exons were most likely lost in the rodent evolution^1^. Along this line, we speculated that some biological processes controlled by human TRPM8 isoforms are shared in mice, but to an unknown extent. In the current study, we reported that the lipidomic fingerprint of TRPM8 isoforms knock-down in LNCaP C4-2b cells shared similarities with the fingerprint of prostate of TRPM8 knock-out mouse with major common contributors in the variance being phosphatidyl choline (PC) and ceramides (**Fig. 4C and 4E**). Indeed, ceramides and PE with long chain FFAs at the sn2 position and very long chain FFAs were expressed at higher levels in all TRPM8 isoforms-silenced LNCaP C4-2b cells and prostates of KOM8 mice (**Tables**). However, PCs with long chain FFAs at the sn2 were expressed at higher level in control LNCaP cells and in KOM8 mice what indicated a clear difference in the modulation of PC homeostasis. We cannot exclude that differences in lipid variations between mouse prostates and LNCaP cells could be due to the prostate tissue heterogeneity. Yet, we must also bear in mind that the silencing of all TRPM8 isoforms in C4-2b LNCaP cells is an acute and probably transient regulation whereas the changes in our KOM8 mouse tissues represent a new stable state selected over different generations of mice. In conclusion, discrepancies between knock-down and knock-out are expected and reveal the direct regulated mechanisms (knock-down) and the adaptative mechanisms (knock-out).

The main common contributors to variance in lipid content in TRPM8(85)-silenced LNCaP cells and in the pool of *wt* mice were SFAs and MUFAs/ PUFAs. This suggests that endogenous variations in TRPM8 control of the SFA/MUFA ratio in the mouse prostate is likely relying on a mechanism similar to the one of LNCaP cells: a TRPM8 form binding AR in order to control *Scd1* gene expression. Although, TRPM8(85) is not expressed in mouse tissues, we cannot exclude that the sole TRPM8 channel plays a similar role than TRPM8 and TRPM8(85) in human. This hypothesis is supported by the variation in FFA content observed in LNCaP cells, where single TRPM8 knock-down induces similar but milder regulations in MUFA content than TRPM8(85) KD. Finally, we cannot exclude that a mouse-specific isoform exists and play the same role as TRPM8(85) in human.

### Lipid droplets accumulation and mitochondrial stress

Silencing of TRPM8(85) led to increased MUFAs, PUFAs and PE content along with a decrease in PC content. These variations in the lipidome were associated with an increase in lipid droplets, mitophagy figures and vacuolization in mitochondria. Lipid droplets size and accumulation have also been shown to depend on lipid metabolism, especially on SFA content^7^ and on the ratio PC/PE ratio (for review see:^8^). More specifically, a decrease in PC/PE ratio can induce an increase in the size of lipid droplets and ER stress but it can also increase mitochondrial respiration and ATP synthesis. Increased SFA, particularly palmitate, has also been reported to trigger ER stress. However, we observed no increase in either ER stress markers or mitochondrial stress markers tested by PCR in TRPM8(85)-silenced cells. This likely suggested that the variations in the lipidome were too mild to trigger stress mechanisms. Nevertheless, we observed lipid droplet accumulation which could have been a consequence of the decrease in PC/PE ratio. However, we cannot exclude the hypothesis that lipid droplets accumulation was the cause of the decrease in PC/PE ratio if PE were more prone to be accumulated in lipid droplets. In conclusion, future studies will be required to investigate the underlying causes and consequences between these two phenomena and their link with TRPM8 isoforms.

## SUPPLEMENTAL FIGURE LEGENDS

**FIGURE S1.**
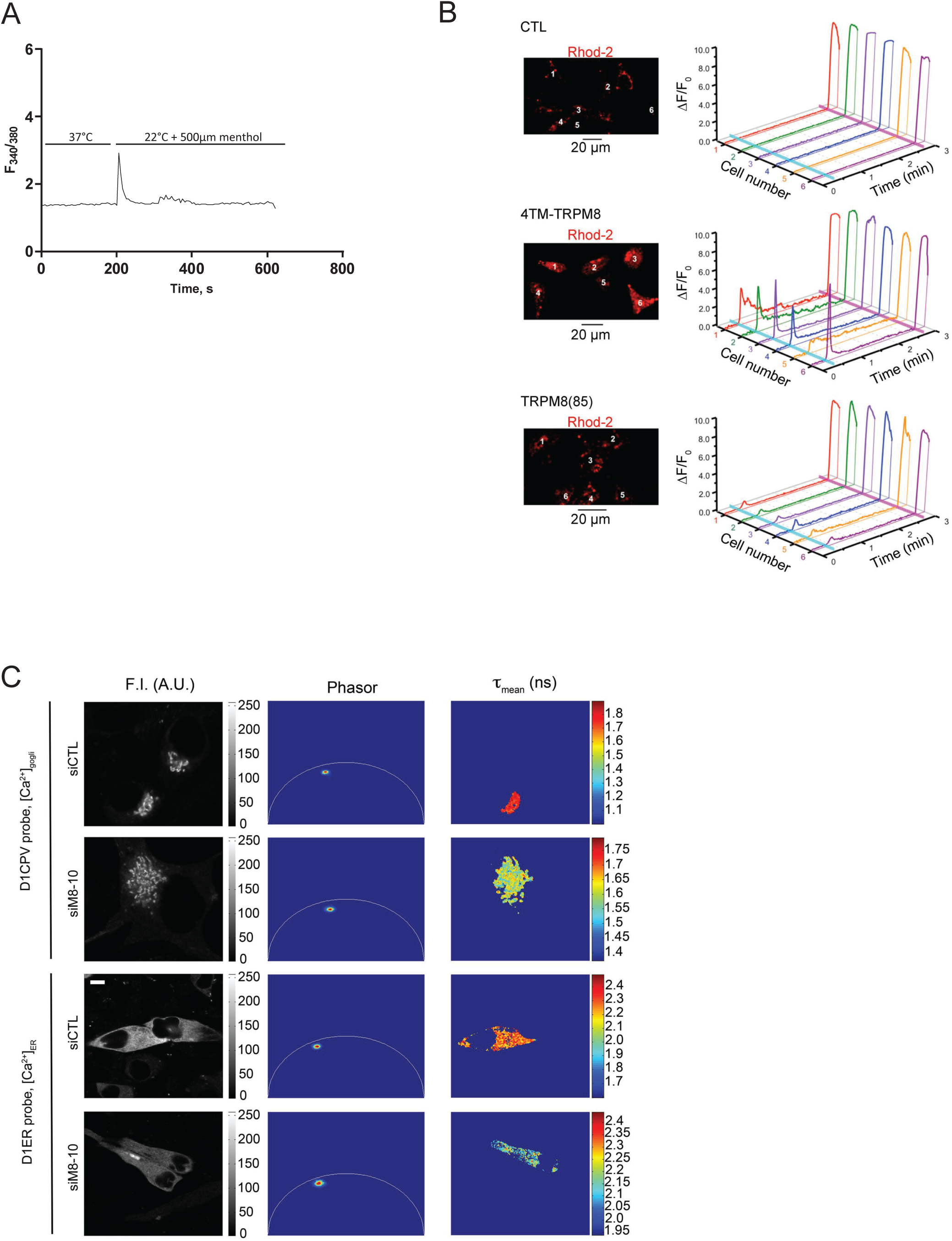
**A,** Changes in the cytosolic Ca^2+^ concentration measured with Fura-2 in HEK cells, transfected with TRPM8(85) encoding vector. The Ca^2+^ signal is elicited by the exposure to cold (22 °C) with 500 μM menthol. **B,** Changes in the mitochondrial Ca^2+^ concentration ([Ca^2+^]_m_) monitored by Rhod2-AM in control HEK cells, transfected with an empty pcDNA4 vector (CTL), HEK cells transfected with 4TM-TRPM8 encoding plasmid or TRPM8(85) encoding vector using x-y time series imaging of rhod-2 fluorescence. The images were acquired at 1.5 Hz from confocal optical slice below 2 μm. The cells were bathed in Ca^2+^-free solution supplemented with 3 mM EGTA and stimulated with 200 μM menthol. To estimate the rhod-2 load, the cells were exposed to 2.5 μM ionomycin at the end of the experiment. The fluorescence intensity (F) was normalized to the averaged fluorescence intensity before menthol application (F_0_). Relative changes in the fluorescence intensity (ΔF/F_0_), averaged within each of 6 cells, denoted by the numbers on the images (left), are plotted over time, respectively (right). Menthol and ionomycin applications are depicted on the 3D plots by vertical cyan and magenta bars, respectively. **C,** Gray-scale coded fluorescence images (left column) were analysed with the phasor plot method (Phasor; 2^nd^ column) to compute FRET-FLIM images (τ_mean_) showing the mean lifetime of the cerulean protein from TCSPC histograms. Intensity in all images is coded as represented by the respective scale bars. Scale bar: 5 µm. The statistical results of these images are presented in the figure 1I.

**FIGURE S2.**
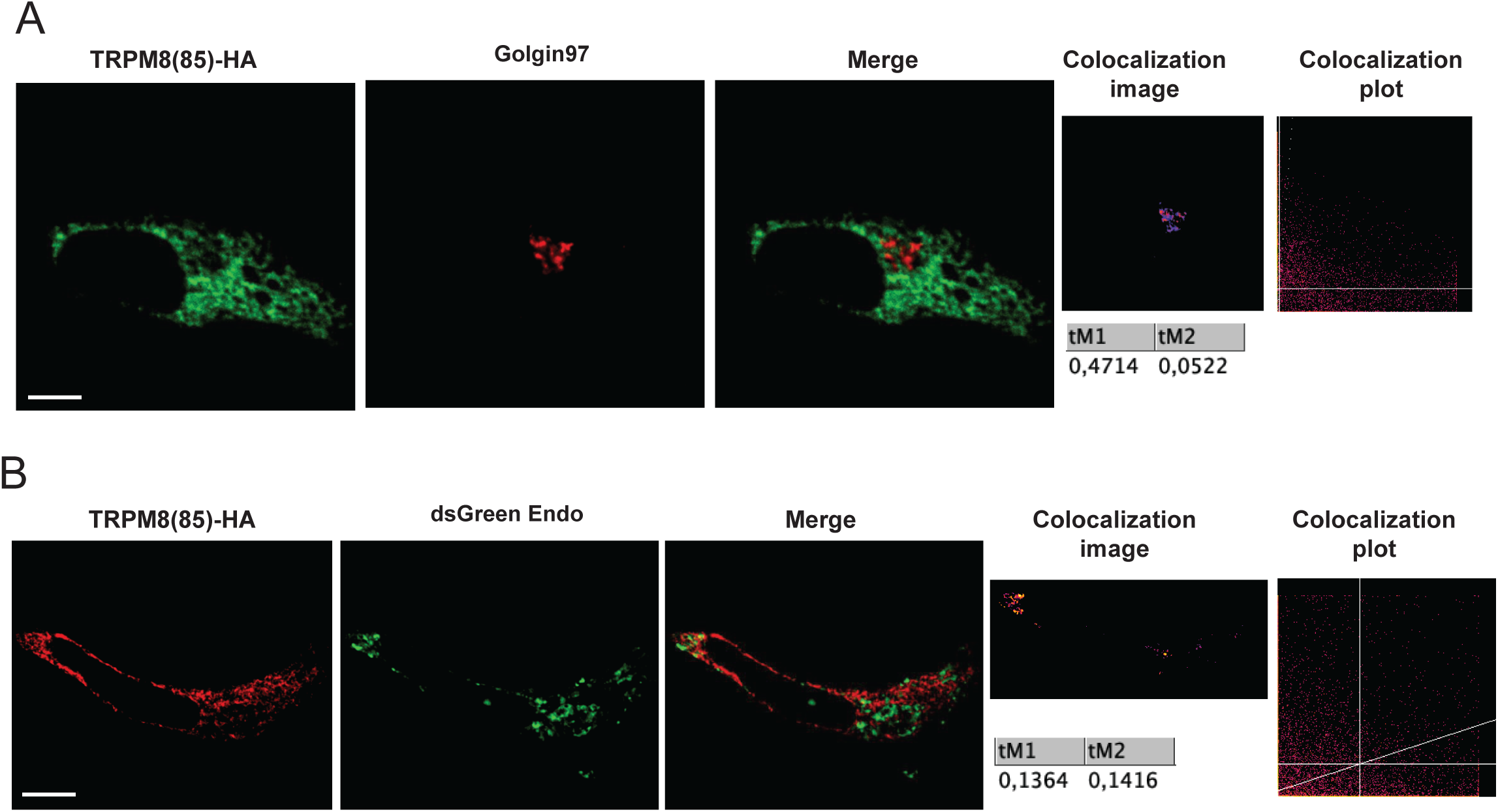
TRPM8(85) is expressed in ER microdomains, partly in the vicinity of GA compartment. Immunofluorescence shows, in green (1^st^ column), the detection of recombinant HA- tagged TRPM8(85) protein in LNCaP C4-2b cells, (2^nd^ column) Golgi marker or endosomal marker and overlay of the two fluorescent signals is shown in the 3^rd^ column (Merge). On the last column, colocalization image, colocalization plot and the results of the colocalization pixel analysis with Manders’ coefficient analysed with ImageJ are presented. Scale bar: 5µm. n=3. **A,** Recombinant HA- tagged TRPM8(85) protein in green with Golgi marker, Golgin-97 in red. **B,** Recombinant HA-tagged TRPM8(85) protein in red with endosomal marker, dsGreen in green.

**FIGURE S3.**
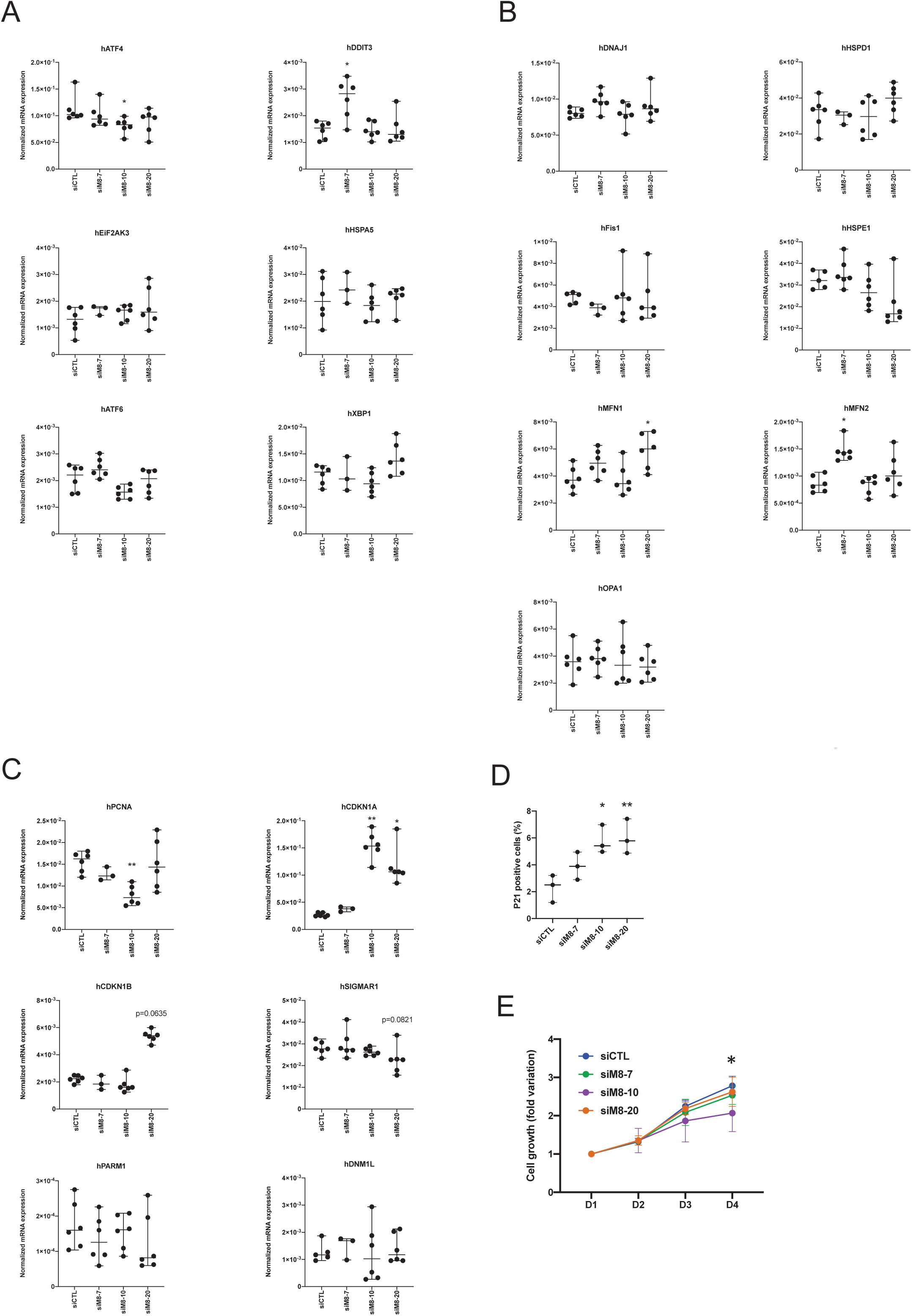
Knock-down of TRPM8(85) increase p21 expression, decrease PCNA and cell growth by 10%. Graph plot showing the normalized expression of transcripts measured by qPCR, in LNCaP C4-2b cells transfected siCTL, siM8-7, siM8-10 and siM8-20 for 3 days. ER stress markers (**A**): hATF4, hDDIT3, hEiF2AK3, hHSPA5, hATF6 and hXBP1. Markers of mitochondrial fusion/fission and stress (**B**): hDNAJ1, hHSPD1, hFis1, hHSPE1, hMFN1, hMFN2 and hOPA1. Markers of cell cycle and cell death (**C**): hPCNA, hCDKN1A, hCDKN1B, hSIGMAR1, hPARM1, and hDNM1L. n=3-6. Unpaired *t* test with Welch’s corrected test was applied and significance reached with p<0.05. **D.** LNCaP C4-2b cells transfected siCTL, siM8-7, siM8-10 and siM8-20 for 3 days were sorted out for p21^CIP1/WAF1^ expression by flow cytometry. Graph plot shows the mean percentage of p21 positive cells in 3 independent experiments. Mean±SD. Unpaired *t* test with Welch’s corrected test was applied and significance reached with p<0.05. **E,** TRPM8(85) knock-down slightly decreases LNCaP C4-2b growth in vitro. Cell growth of LNCaP C4-2b cells transfected with siCTL, siM8-7 or siM8-10 was measured with CellTiter 96® AQueous Non-Radioactive Cell Proliferation Assay (Promega), each day for 3 days of culture. Values were normalized to value at D1. Experiments were performed three times independently. Graph plot shows Mean±SD. One-way ANOVA with paired test over time was done.

**FIGURE S4.**
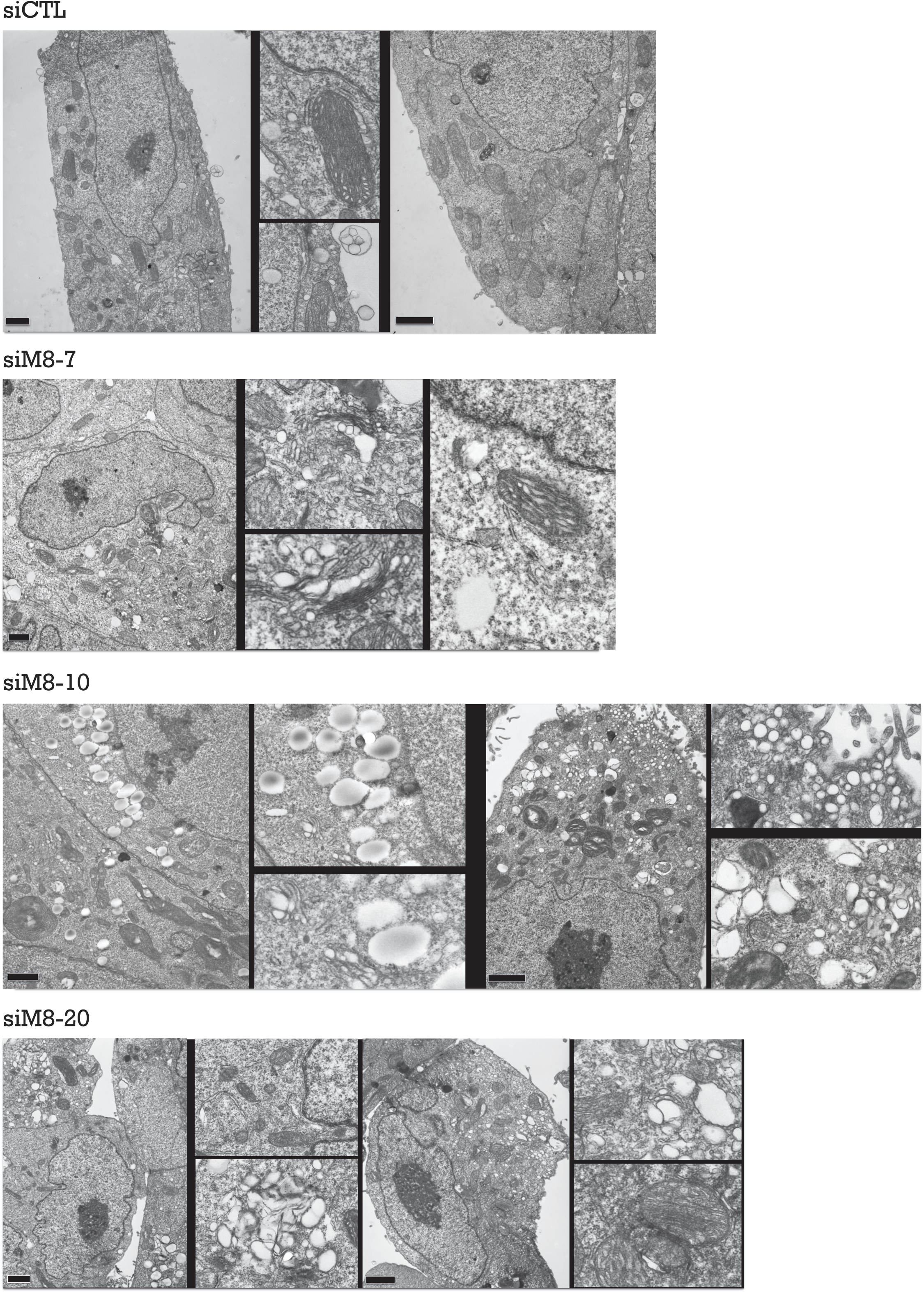
TRPM8(85) knock-down induced GA expansion and fragmentation in small vesicles, and is associated with accumulation of lipid droplet and secretion vesicles. Representative TEM micrograph shows cells transfected with siCTL, siM8-7, siM8-10 or siM8-20 for 72h. Scale bars: 1 µm.

**FIGURE S5.**
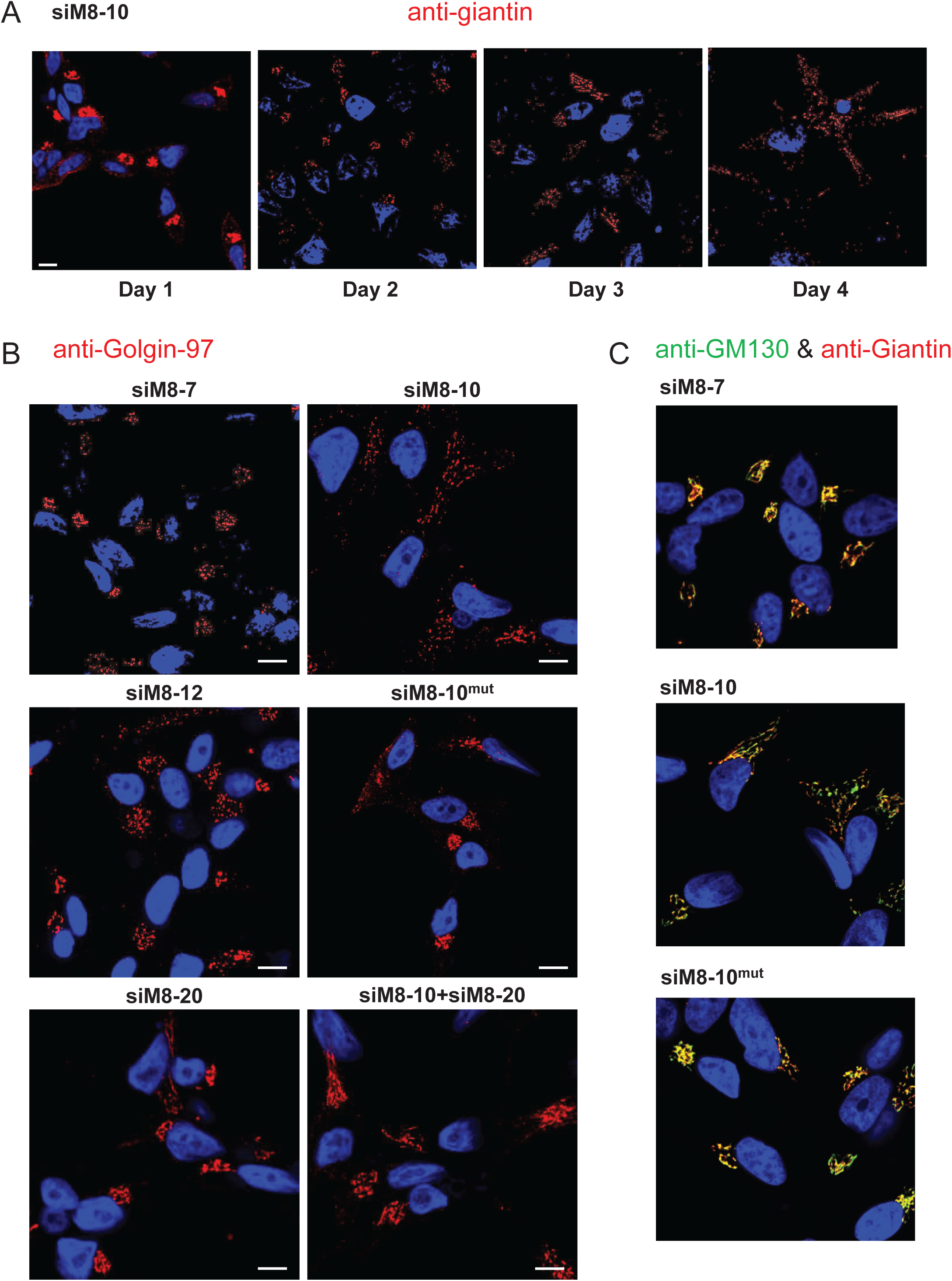
A 4-day TRPM8(85) knock-down induces GA (cis/medial-Golgi and TGN) expansion in LNCaP C4-2b cells and required expression of 4TM-TRPM8. **A.** Immunofluorescent labelling of Giantin in LNCaP C4-2b cells shows the modification of the cis-Golgi morphology over 96h of TRPM8(85) knocking-down (siM8-10). **B.** Immunofluorescent labelling of Golgin97 in LNCaP C4-2b cells shows the modification in the TGN morphology after a 96h-knocking down of full length TRPM8 (siM8-7), TRPM8(85)+TRPM8 (siM8-10 and siM8-12), mutated siM8-10 sequence (siM8- 10^mut^) used as a negative control, concomitant knocking down of TRPM8, TRPM8(85) and 4TM- TRPM8 (siM8-20), concomitant knocking down of with siM8-10 and siM8-20. Scale bar: 10 µm. **C.** Immunofluorescent labelling of Giantin (cis/medial-Golgi) and GM130 (cis-Golgi) in LNCaP C4-2b cells shows the modification of the cis-Golgi morphology over 96h after knocking down with either siM8-7, siM8-10 and mutated siM8-10 sequence (siM8-10^mut^). Scale bar: 10 µm.

**FIGURE S6.**
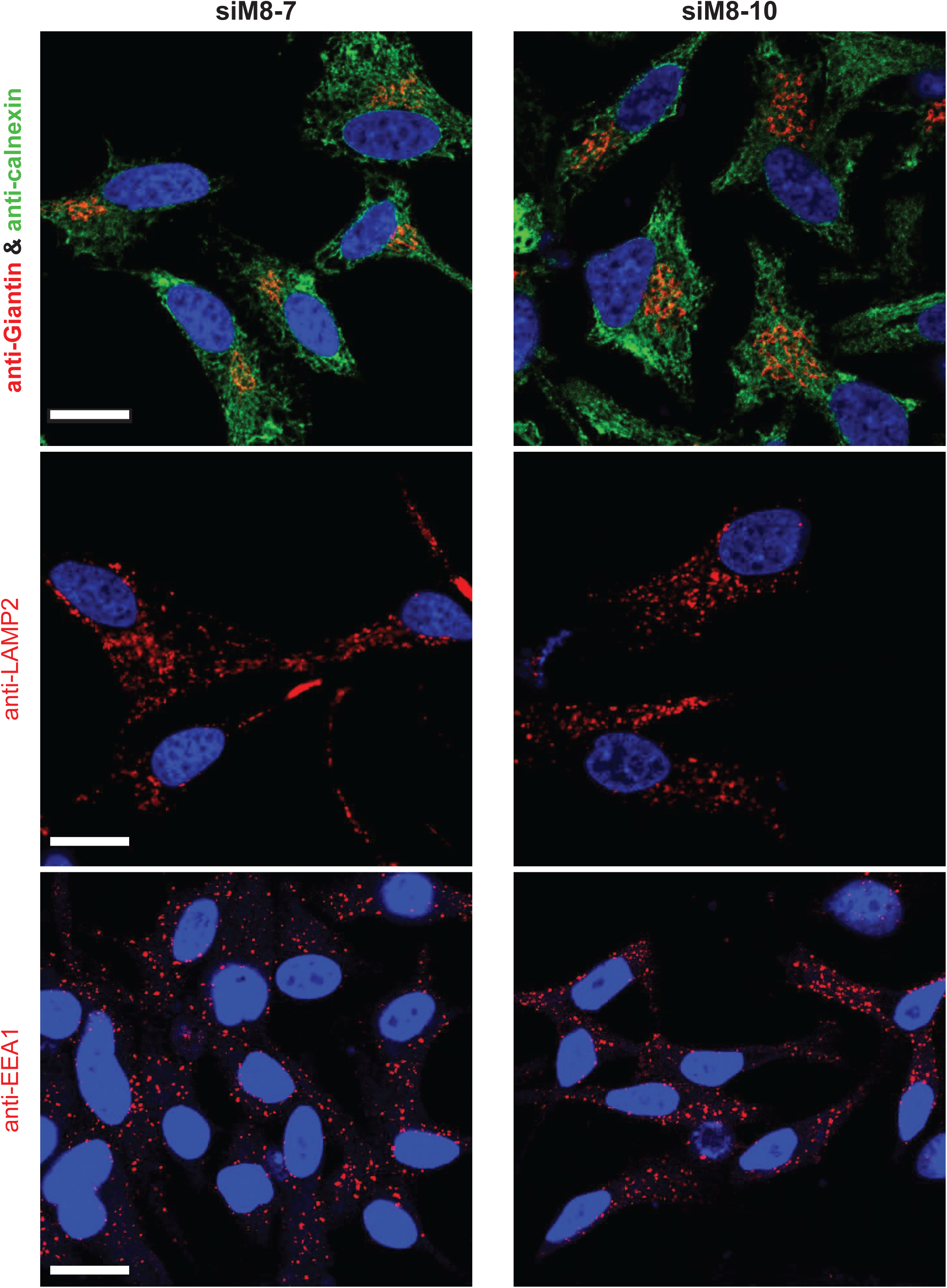
TRPM8(85) knock-down does not phenotypically modifies ER structure, lysosome content and endosome content in prostate cancer cells. Immunofluorescent labelling of Giantin (*cis/medial*-Golgi marker), Calnexin (ER marker), LAMP2 (lysosome marker) and EEA1 (endosome marker) in LNCaP C4-2b cells transfected either with siM8-7 or siM8-10 for 96h. Dapi staining labelled nuclei. Scale bar: 10 µm.

**FIGURE S7.**
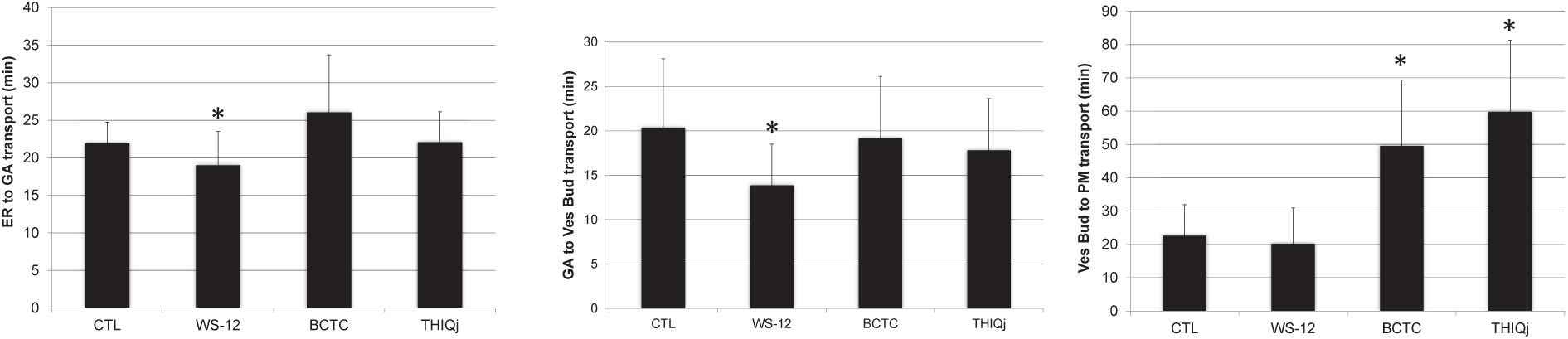
Altered cargo transport upon targeting TRPM8 in LNCaP cells. VSVG-GFP assay used to monitor the transport time of protein from ER to plasmalemma. The three plots show the time (min) of trafficking from ER to Golgi apparatus (GA), Golgi apparatus (GA) to vesicle budding (Ves Bud) and vesicle budding (Ves Bud) to plasmalemma (PM). LNCaP C4-2b cells were treated with a TRPM8 agonist (5µM WS-12) or two TRPM8 antagonists (10µM BCTC and 25µM THIQj) at the onset of the recording. Data are presented as Mean±SD. T-tests were performed against the control group with p value< 0.05 considered significant. n=3 independent experiments were performed.

**FIGURE S8.**
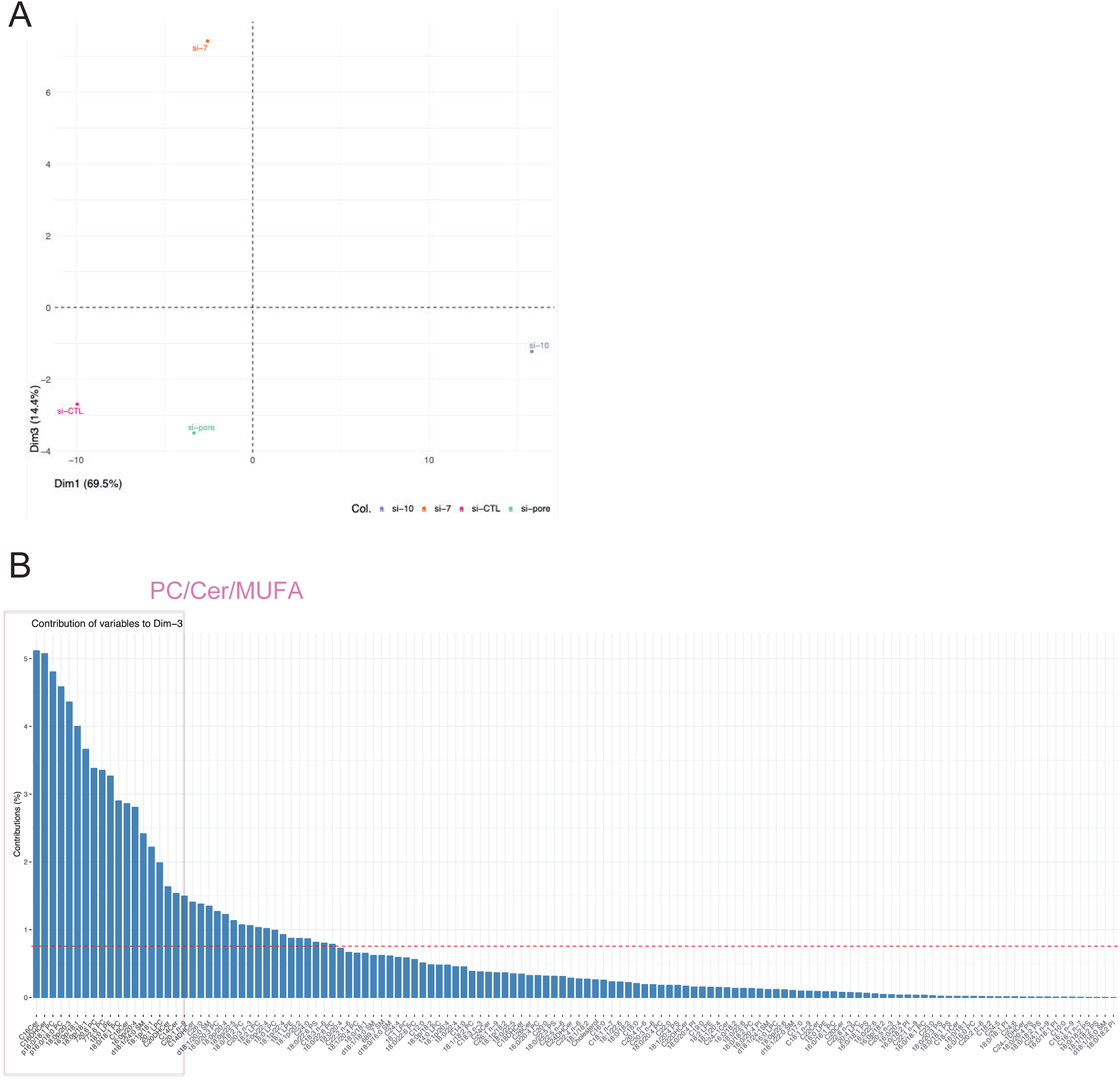
**A.** Principle Component Analysis (PCA) biplot depicting the relationship between the lipid species and the siRNA treatment in LNCaP C4-2b cells. The first and third dimension (dim1 and dim3, respectively) were used to scared the siM8-7 and siM8-10 groups. Each of the four points represents the average lipid profile for one siRNA treatment. **B.** The histogram shows the main contributors among the lipid species in the dim3 of the PCA. The pink square highlights the top ranking of these contributors and is mainly composed of phosphatidylcholine (PC), ceramides (Cer) and monounsaturated fatty acids (MUFAs).

**FIGURE S9.**
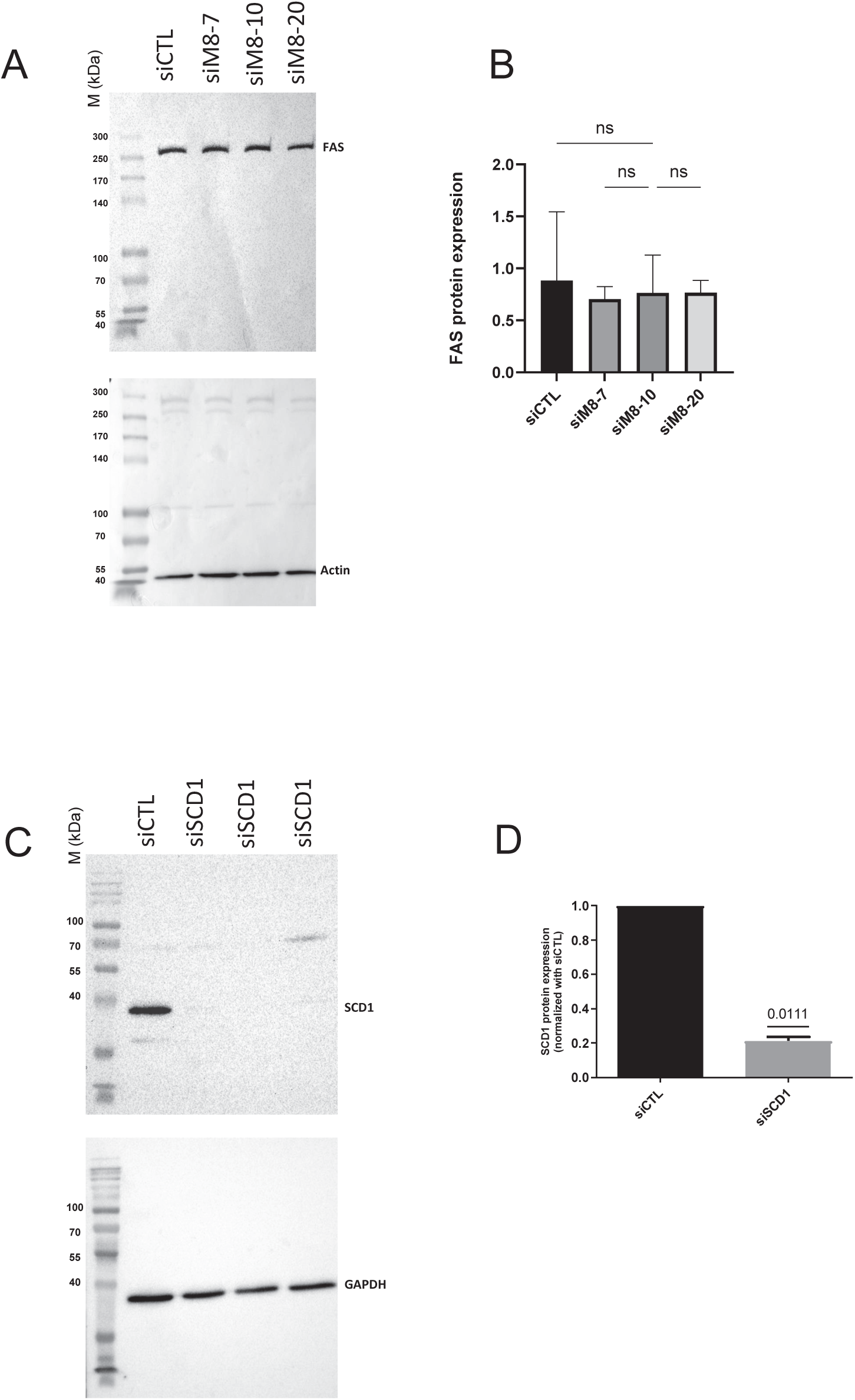
Fatty acid synthase (FAS) protein in unchanged in TRPM8(85)-silenced LNCaP cells. **A**. Western blot shows detection of the FAS protein expression in cells transfected with the different siRNAs targeting TRPM8 and its isoforms. The house keeping gene shown is actin. Quantitation of FAS protein expression. Data are presented as Median±95%CI. Kruskal Wallis test was performed. **B.** SCD1 protein expression was assessed by western blot upon its knocking down using siSCD1. (A) Western blot presentation with siCTL and siSCD1. The house keeping gene used is GAPDH. (B) Quantitation of SCD1 protein expression in both siCTL and siSCD1 (data normalized to siCTL). Data are presented as Mean±SEM. One sample t test was performed with p value< 0.05.

**FIGURE S10.**
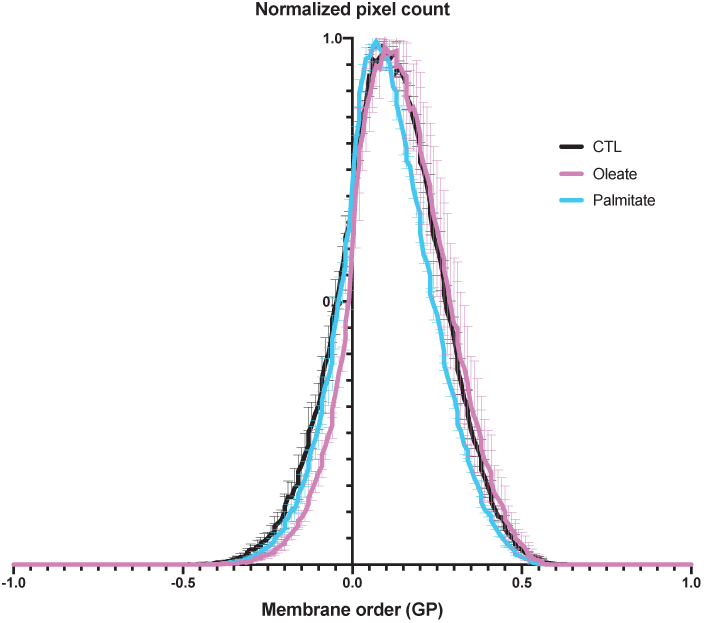
Membrane order analysis using Laurdan (6-Dodecanoyl-2- dimethylaminonaphthalene) in LNCaP cells following lipid treatment for 3 days. Generalised polarisation (GP) values for LNCaP cells after 3 days treatment with 50 µM of either palmitate (SFA) or oleate (MUFA) are presented. GP is calculated from the fluorescence spectrum of Laurdan.

## SUPPLEMENTAL TABLE

**Supplemental table 1:**
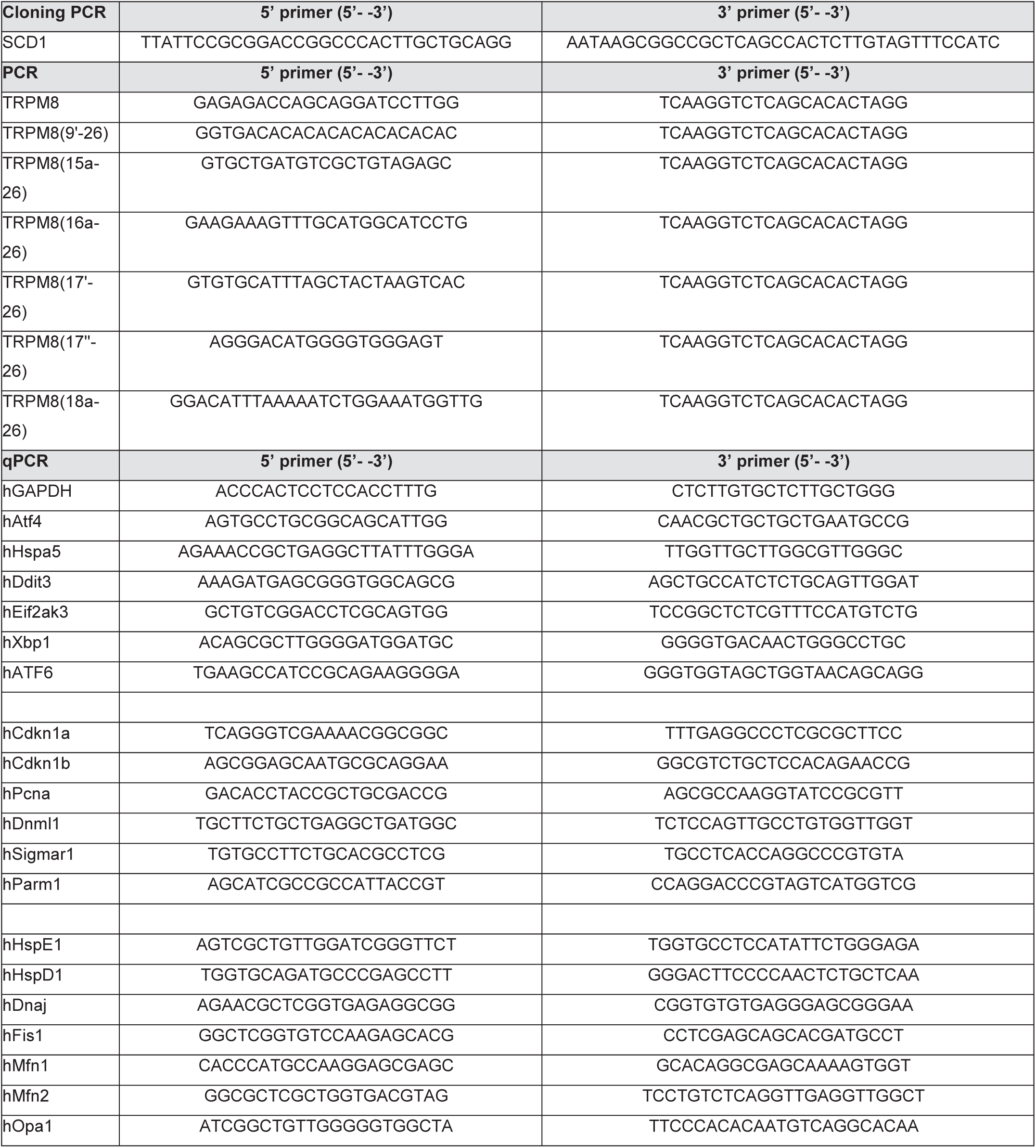

## REFERENCES

1. Stein, R. J. et al. Cool (TRPM8) and hot (TRPV1) receptors in the bladder and male genital tract. J. Urol. 172, 1175–1178 (2004).

2. Tsavaler, L., Shapero, M. H., Morkowski, S. & Laus, R. Trp-p8, a novel prostate-specific gene, is up- regulated in prostate cancer and other malignancies and shares high homology with transient receptor potential calcium channel proteins. Cancer Res 61, 3760–9 (2001).

3. McKemy, D. D., Neuhausser, W. M. & Julius, D. Identification of a cold receptor reveals a general role for TRP channels in thermosensation. Nature 416, 52–8 (2002).

4. Peier, A. M. et al. A TRP channel that senses cold stimuli and menthol. Cell 108, 705–15 (2002).

5. Bautista, D. M. et al. The menthol receptor TRPM8 is the principal detector of environmental cold. Nature 448, 204–8 (2007).

6. Dhaka, A. et al. TRPM8 is required for cold sensation in mice. Neuron 54, 371–8 (2007).

7. Colburn, R. W. et al. Attenuated cold sensitivity in TRPM8 null mice. Neuron 54, 379–386 (2007).

8. Liu, Y. et al. TRPM8 channels: A review of distribution and clinical role. Eur. J. Pharmacol. 882, 173312 (2020).

9. Denda, M., Tsutsumi, M. & Denda, S. Topical application of TRPM8 agonists accelerates skin permeability barrier recovery and reduces epidermal proliferation induced by barrier insult: role of cold- sensitive TRP receptors in epidermal permeability barrier homoeostasis. Exp. Dermatol. 19, 791–795 (2010).

10. Bidaux, G., Borowiec, A. S., Prevarskaya, N. & Gordienko, D. Fine-tuning of eTRPM8 expression and activity conditions keratinocyte fate. Channels Austin 10, 320–31 (2016).

11. Bidaux, G. et al. Epidermal TRPM8 channel isoform controls the balance between keratinocyte proliferation and differentiation in a cold-dependent manner. Proc Natl Acad Sci U A 112, E3345–54 (2015).

12. De Blas, G. A. et al. TRPM8, a versatile channel in human sperm. PloS One 4, e6095 (2009).

13. Borowiec, A. S. et al. Cold/menthol TRPM8 receptors initiate the cold-shock response and protect germ cells from cold-shock-induced oxidation. FASEB J 30, 3155–70 (2016).

14. Zhang, L. & Barritt, G. J. Evidence that TRPM8 is an androgen-dependent Ca2+ channel required for the survival of prostate cancer cells. Cancer Res 64, 8365–73 (2004).

15. Thebault, S. et al. Novel role of cold/menthol-sensitive transient receptor potential melastatine family member 8 (TRPM8) in the activation of store-operated channels in LNCaP human prostate cancer epithelial cells. J Biol Chem 280, 39423–35 (2005).

16. Bidaux, G. et al. Prostate cell differentiation status determines transient receptor potential melastatin member 8 channel subcellular localization and function. J Clin Invest 117, 1647–57 (2007).

17. Bidaux, G. et al. Evidence for specific TRPM8 expression in human prostate secretory epithelial cells: functional androgen receptor requirement. Endocr Relat Cancer 12, 367–82 (2005).

18. Fuessel, S. et al. Multiple tumor marker analyses (PSA, hK2, PSCA, trp-p8) in primary prostate cancers using quantitative RT-PCR. Int J Oncol **23**, 221–8 (2003).

19. Kiessling, A. et al. Identification of an HLA-A*0201-restricted T-cell epitope derived from the prostate cancer-associated protein trp-p8. Prostate 56, 270–9 (2003).

20. Gkika, D., Flourakis, M., Lemonnier, L. & Prevarskaya, N. PSA reduces prostate cancer cell motility by stimulating TRPM8 activity and plasma membrane expression. Oncogene 29, 4611–6 (2010).

21. Henshall, S. M. et al. Survival analysis of genome-wide gene expression profiles of prostate cancers identifies new prognostic targets of disease relapse. Cancer Res 63, 4196–203 (2003).

22. Blanquart, S. et al. Evolution of the human cold/menthol receptor, TRPM8. Mol. Phylogenet. Evol. **136**, 104–118 (2019).

23. Bidaux, G. et al. Regulation of activity of transient receptor potential melastatin 8 (TRPM8) channel by its short isoforms. J Biol Chem 287, 2948–62 (2012).

24. Bidaux, G. et al. Targeting of short TRPM8 isoforms induces 4TM-TRPM8-dependent apoptosis in prostate cancer cells. Oncotarget 7, 29063–80 (2016).

25. Fernandez, J. A. et al. Short Isoforms of the Cold Receptor TRPM8 Inhibit Channel Gating by Mimicking Heat Action rather than Chemical Inhibitors. J Biol Chem (to be published in the same volume),.

26. Bidaux, G. et al. 4TM-TRPM8 channels are new gatekeepers of the ER-mitochondria Ca(2+) transfer. Biochim Biophys Acta Mol Cell Res 1865, 981–994 (2018).

27. de Mendoza, D. Temperature sensing by membranes. Annu. Rev. Microbiol. 68, 101–116 (2014).

28. Inda, M. E. et al. A lipid-mediated conformational switch modulates the thermosensing activity of DesK. Proc. Natl. Acad. Sci. U. S. A. 111, 3579–3584 (2014).

29. Saita, E. et al. A coiled coil switch mediates cold sensing by the thermosensory protein DesK. Mol. Microbiol. 98, 258–271 (2015).

30. Brauchi, S., Orta, G., Salazar, M., Rosenmann, E. & Latorre, R. A hot-sensing cold receptor: C-terminal domain determines thermosensation in transient receptor potential channels. J. Neurosci. Off. J. Soc. Neurosci. 26, 4835–4840 (2006).

31. Tsuruda, P. R., Julius, D. & Minor, D. L. Coiled coils direct assembly of a cold-activated TRP channel. Neuron 51, 201–212 (2006).

32. Erler, I. et al. Trafficking and assembly of the cold-sensitive TRPM8 channel. J. Biol. Chem. 281, 38396–38404 (2006).

33. Rohács, T., Lopes, C. M. B., Michailidis, I. & Logothetis, D. E. PI(4,5)P2 regulates the activation and desensitization of TRPM8 channels through the TRP domain. Nat. Neurosci. 8, 626–634 (2005).

34. Liu, B. & Qin, F. Functional control of cold- and menthol-sensitive TRPM8 ion channels by phosphatidylinositol 4,5-bisphosphate. J. Neurosci. Off. J. Soc. Neurosci. 25, 1674–1681 (2005).

35. Vanden Abeele, F., et al. Ca2+-independent phospholipase A2-dependent gating of TRPM8 by lysophospholipids. J Biol Chem 281, 40174–82 (2006).

36. Andersson, D. A., Nash, M. & Bevan, S. Modulation of the cold-activated channel TRPM8 by lysophospholipids and polyunsaturated fatty acids. J Neurosci 27, 3347–55 (2007).

37. Bidaux, G. et al. FRET Image Correlation Spectroscopy Reveals RNAPII-Independent P-TEFb Recruitment on Chromatin. Biophys J 114, 522–533 (2018).

38. Sanchez, S. A., Tricerri, M. A. & Gratton, E. Laurdan generalized polarization fluctuations measures membrane packing micro-heterogeneity in vivo. Proc. Natl. Acad. Sci. U. S. A. 109, 7314–7319 (2012).

39. Grynkiewicz, G., Poenie, M. & Tsien, R. Y. A new generation of Ca2+ indicators with greatly improved fluorescence properties. J Biol Chem 260, 3440–50 (1985).

40. Chouabe, C. et al. Reduction of I(Ca,L) and I(to1) density in hypertrophied right ventricular cells by simulated high altitude in adult rats. J. Mol. Cell. Cardiol. 29, 193–206 (1997).

41. Vial, G. et al. Imeglimin normalizes glucose tolerance and insulin sensitivity and improves mitochondrial function in liver of a high-fat, high-sucrose diet mice model. Diabetes 64, 2254–2264 (2015).

42. Tran, T. T. T. et al. Short Term Palmitate Supply Impairs Intestinal Insulin Signaling via Ceramide Production. J. Biol. Chem. 291, 16328–16338 (2016).

43. Blondelle, J., Pais de Barros, J.-P., Pilot-Storck, F. & Tiret, L. Targeted Lipidomic Analysis of Myoblasts by GC-MS and LC-MS/MS. Methods Mol. Biol. Clifton NJ 1668, 39–60 (2017).

44. Shaner, R. L. et al. Quantitative analysis of sphingolipids for lipidomics using triple quadrupole and quadrupole linear ion trap mass spectrometers. J. Lipid Res. 50, 1692–1707 (2009).

45. Retra, K. et al. A simple and universal method for the separation and identification of phospholipid molecular species. Rapid Commun. Mass Spectrom. RCM 22, 1853–1862 (2008).

46. Phelps, C. B. & Gaudet, R. The role of the N terminus and transmembrane domain of TRPM8 in channel localization and tetramerization. J Biol Chem 282, 36474–80 (2007).

47. Lissandron, V., Podini, P., Pizzo, P. & Pozzan, T. Unique characteristics of Ca2+ homeostasis of the trans-Golgi compartment. Proc. Natl. Acad. Sci. U. S. A. 107, 9198–9203 (2010).

48. Hirschberg, K. et al. Kinetic analysis of secretory protein traffic and characterization of golgi to plasma membrane transport intermediates in living cells. J. Cell Biol. 143, 1485–1503 (1998).

49. Sweeney, D. A., Siddhanta, A. & Shields, D. Fragmentation and re-assembly of the Golgi apparatus in vitro. A requirement for phosphatidic acid and phosphatidylinositol 4,5-bisphosphate synthesis. J. Biol. Chem. **277**, 3030–3039 (2002).

50. Enoch, H. G., Catalá, A. & Strittmatter, P. Mechanism of rat liver microsomal stearyl-CoA desaturase. Studies of the substrate specificity, enzyme-substrate interactions, and the function of lipid. J. Biol. Chem. 251, 5095–5103 (1976).

51. Barelli, H. & Antonny, B. Lipid unsaturation and organelle dynamics. Curr. Opin. Cell Biol. 41, 25–32 (2016).

52. Audet-Walsh, É. et al. SREBF1 Activity Is Regulated by an AR/mTOR Nuclear Axis in Prostate Cancer. Mol. Cancer Res. MCR 16, 1396–1405 (2018).

53. Grolez, G. P. et al. TRPM8-androgen receptor association within lipid rafts promotes prostate cancer cell migration. Cell Death Dis. 10, 652 (2019).

54. Gkika, D. et al. Testosterone-androgen receptor: The steroid link inhibiting TRPM8-mediated cold sensitivity. FASEB J. Off. Publ. Fed. Am. Soc. Exp. Biol. 34, 7483–7499 (2020).

55. Ramazanov, B. R. et al. Calcium flow at ER-TGN contact sites facilitates secretory cargo export. Mol. Biol. Cell 35, ar50 (2024).

56. Ford, C., Parchure, A., von Blume, J. & Burd, C. G. Cargo sorting at the trans-Golgi network at a glance. J. Cell Sci. 134, jcs259110 (2021).

57. Mikhaylova, M. et al. Calneurons provide a calcium threshold for trans-Golgi network to plasma membrane trafficking. Proc. Natl. Acad. Sci. U. S. A. 106, 9093–9098 (2009).

58. Clermont, Y., Rambourg, A. & Hermo, L. Trans-Golgi network (TGN) of different cell types: three- dimensional structural characteristics and variability. Anat. Rec. 242, 289–301 (1995).

59. Sens, P. & Rao, M. (Re)modeling the Golgi. Methods Cell Biol. 118, 299–310 (2013).

60. Sarmento, M. J. et al. The expanding organelle lipidomes: current knowledge and challenges. Cell. Mol. Life Sci. CMLS 80, 237 (2023).

61. Lira, R. B. et al. The underlying mechanical properties of membranes tune their ability to fuse. J. Biol. Chem. 299, 105430 (2023).

62. Lita, A. et al. IDH1 mutations induce organelle defects via dysregulated phospholipids. Nat. Commun. 12, 614 (2021).

63. Payet, L.-A. et al. Saturated fatty acids alter the late secretory pathway by modulating membrane properties. Traffic Cph. Den. 14, 1228–1241 (2013).

64. Rog, T. & Koivuniemi, A. The biophysical properties of ethanolamine plasmalogens revealed by atomistic molecular dynamics simulations. Biochim. Biophys. Acta 1858, 97–103 (2016).

65. Vanni, S. et al. Use of biomarkers in triage of patients with suspected stroke. J. Emerg. Med. 40, 499– 505 (2011).

66. Asuthkar, S. et al. The TRPM8 protein is a testosterone receptor: II. Functional evidence for an ionotropic effect of testosterone on TRPM8. J. Biol. Chem. **290**, 2670–2688 (2015).

67. Asuthkar, S. et al. The TRPM8 protein is a testosterone receptor: I. Biochemical evidence for direct TRPM8-testosterone interactions. J. Biol. Chem. **290**, 2659–2669 (2015).

68. Marklund, M. et al. Spatio-temporal analysis of prostate tumors in situ suggests pre-existence of treatment-resistant clones. Nat. Commun. 13, 5475 (2022).

## References

1. Blanquart, S. et al. Evolution of the human cold/menthol receptor, TRPM8. Mol. Phylogenet. Evol. **136**, 104–118 (2019).

2. Bidaux, G. et al. Epidermal TRPM8 channel isoform controls the balance between keratinocyte proliferation and differentiation in a cold-dependent manner. Proc Natl Acad Sci U A 112, E3345–54 (2015).

3. Bidaux, G. et al. Targeting of short TRPM8 isoforms induces 4TM-TRPM8-dependent apoptosis in prostate cancer cells. Oncotarget 7, 29063–80 (2016).

4. Bidaux, G. et al. 4TM-TRPM8 channels are new gatekeepers of the ER-mitochondria Ca(2+) transfer. Biochim Biophys Acta Mol Cell Res 1865, 981–994 (2018).

5. Borowiec, A. S. et al. Cold/menthol TRPM8 receptors initiate the cold-shock response and protect germ cells from cold-shock-induced oxidation. FASEB J 30, 3155–70 (2016).

6. Saito, S. & Shingai, R. Evolution of thermoTRP ion channel homologs in vertebrates. Physiol Genomics 27, 219–30 (2006).

7. Thomas, P. et al. Differential routing and disposition of the long-chain saturated fatty acid palmitate in rodent vs human beta-cells. Nutr. Diabetes 12, 22 (2022).

8. van der Veen, J. N. et al. The critical role of phosphatidylcholine and phosphatidylethanolamine metabolism in health and disease. Biochim. Biophys. Acta Biomembr. 1859, 1558–1572 (2017).

